# Model-assisted high-throughput phenotyping of photosynthetic acclimation

**DOI:** 10.64898/2026.09.21.753097

**Authors:** Emilio Villar Alegria, Nicholas J. Russell, Tsu-Wei Chen

## Abstract

Photosynthesis is a key determinant of crop productivity, yet conventional gas-exchange measurements are labor-intensive and unsuitable for high-throughput assessment of dynamic photosynthetic acclimation across large genotype panels. Consequently, genetic variation in photosynthetic acclimation remains poorly characterized and difficult to exploit in breeding programs. Here we present a computational pipeline that integrates rapid optical sensing (chlorophyll meters and hyperspectral reflectance) with a mechanistic model of photosynthetic protein-turnover to estimate dynamic nitrogen allocation among photosynthetic components. The pipeline was demonstrated using 60 winter wheat (*Triticum aestivum* L.) cultivars sampled at 10 time points spanning leaf emergence to senescence. Uncertainties associated with each pipeline component were quantified and benchmarked against gas-exchange ground-truth measurements under three controlled-environment light and temperature regimes. Dynamic nitrogen allocation to light harvesting, electron transport, and carboxylation was characterized using three biologically interpretable parameters: maximum synthesis rate, degradation rate, and age-dependent decline in synthesis. Although hyperspectral models showed moderate predictive accuracy for photosynthetic capacity (R² ≈ 0.49), prediction errors were predominantly random, allowing robust parameter estimation when observations across the leaf lifespan were integrated. The pipeline successfully resolved genotype-by-environment interactions in photosynthetic acclimation, with environmental and interaction effects contributing more strongly than genotype main effects to nitrogen dynamics. These results demonstrate a scalable, uncertainty-aware sensor-to-trait architecture for dynamic physiological phenotyping that enables high-throughput germplasm screening across diverse environments and provides novel physiological selection targets for improving crop adaptation under variable climates.

## 1. Introduction

Photosynthesis sustains terrestrial productivity through atmospheric CO₂ fixation (Long et al., 2006; Zhu et al., 2010), but its performance also depends on how leaves allocate a finite nitrogen budget among three functional domains: light harvesting by pigment-protein antenna complexes, electron transport through the thylakoid membranes to generate ATP and NADPH, and carboxylation by Rubisco in the Calvin-Benson cycle (Nelson and Junge, 2015; Niyogi and Truong, 2013). Together, these processes contain most of the nitrogen in green leaves and compete for this limited resource (Evans and Clarke, 2019; Hikosaka, 2014; Niinemets, 2023). Optimal nitrogen partitioning should minimise co-limitation among them, yet allocation strategies vary substantially among species and cultivars (Buckley et al., 2013; Hikosaka et al., 2016; Onoda et al., 2017). Because nitrogen fertilisation is both a major agricultural input and a source of environmental costs, understanding the dynamics of photosynthetic nitrogen (*N*_ph_) could help improve nitrogen use while reducing fertiliser demand and its associated impacts (Martre et al., 2024; Masclaux-Daubresse et al., 2010).

Those nitrogen investments are not fixed. Photosynthetic acclimation, defined here as the dynamic adjustment of nitrogen allocation among photosynthetic components, operates across multiple timescales and remains an underexploited target for crop improvement (Calzadilla and Johnson, 2025). Short-term responses involve enzyme activation within minutes (Acevedo-Siaca et al., 2020; Gao et al., 2024; Kaiser et al., 2015), whereas longer-term acclimation involves reorganisation of thylakoid proteins and rebalancing of nitrogen between light harvesting and carbon metabolism (Athanasiou et al., 2010; Gjindali and Johnson, 2023; Townsend et al., 2018; Walters, 2005). Genotypes differ in these acclimation responses (Pao et al., 2023; Salter et al., 2020), making them potentially relevant for wheat, which provides approximately 20% of global calories but has annual yield gains below those required to meet projected demand by 2050 (Dixon et al., 2009; Voss-Fels et al., 2019). However, acclimation remains poorly integrated into breeding because dynamic physiological responses are difficult to phenotype across diverse germplasm and environments (Cutolo et al., 2023; Fu et al., 2022; Ort et al., 2015; Taylor and Long, 2019).

This limitation is primarily methodological. Gas exchange is the reference method for measuring photosynthetic capacity, but the CO_2_-response curves (*A/C*_i_ curves) and light curves used to estimate it can require up to one hour per leaf, strict environmental control, and careful handling of several technical sources of error (Bellasio et al., 2016; Busch et al., 2024; Flexas et al., 2007; Moualeu-Ngangue et al., 2017). Rapid and comparatively accessible measurements provide an alternative. SPAD measurements estimate chlorophyll content within seconds (Parry et al., 2014), while leaf hyperspectral reflectance can be combined with statistical models to predict photosynthetic capacity parameters such as maximum electron transport (*J*_max_) and maximum carboxylation capacity (*V*_cmax_) (Burnett et al., 2021; Fu et al., 2019; Lamour et al., 2026; Silva-Perez et al., 2018). These estimates can then be converted into functional nitrogen pools using established relationships between photosynthetic proteins and catalytic activity (Buckley et al., 2013; Pao et al., 2019a, 2020).

The value of these rapid measurements is not only their throughput, but also that their accuracy enables to capture aging and acclimation across leaf lifespan. Acclimation is a temporal process and therefore requires repeated observations as a leaf develops (Gjindali and Johnson, 2023), rather than a single measurement. The speed of SPAD and hyperspectral sensing, typically less than one minute per measurement, makes these trajectories feasible across many genotypes and provides the temporal resolution needed to fit mechanistic models of photosynthetic acclimation. For example, a turnover model of photosynthetic protein describes changes in three photosynthetic nitrogen pools—light harvesting (*N*_c_), electron transport (*N*_j_), and carboxylation (*N*_v_)—through differential equations that balance synthesis and degradation (Pao et al., 2019a). Their sum, *N*_ph_, represents nitrogen invested in photosynthetic functions and excludes structural, storage, and other metabolic nitrogen. The model describes the rate of change in a nitrogen pool *x* (*N*_x_, where *x* is either *c*, *j*, or *v*) as:

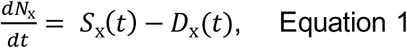

where *S*_x_(*t*) and *D*_x_(*t*) represent instantaneous synthesis and degradation rates (mmol N m⁻² °Cd⁻¹) at leaf age *t* (thermal time, °Cd). Protein degradation follows first-order kinetics, proportional to the existing pool size *N*_x_ (Pao et al., 2019a):

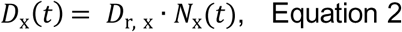

where *D*_r,x_ is the pool-specific degradation constant (°Cd⁻¹). Synthesis capacity declines with leaf age following a logistic function:

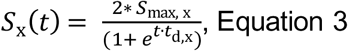

where *S*_max_,_x_ is the maximum synthesis rate during early leaf development and *t*_d_,_x_ describes the age-related decline in synthesis. Together with *D*_r_,_x_, these parameters provide biologically interpretable measures of acclimation: *S*_max_ describes protein accumulation capacity, *t*_d_ describes how long synthesis is sustained as the leaf ages, and *D*_r_ describes protein turnover. Fitting the model to temporal trajectories of *N*_c_, *N*_j_, and *N*_v_ therefore allows nitrogen allocation and turnover to be compared among genotypes and environments (Pao et al., 2023, 2019b).

In this study, we 1) present a computational and data-processing pipeline (Fig. 1, for details, see Fig. S1) that integrates rapid optical sensing (chlorophyll meters and hyperspectral reflectance) with the model presented above to open photosynthetic acclimation to high-throughput phenotyping; 2) quantified and benchmarked errors and uncertainties associated with each pipeline component against gas-exchange ground-truth measurements; 3) demonstrate the pipeline using 60 wheat cultivars, including 50 cultivars representing six decades of German breeding and 10 of exotic origin, grown under constant conditions (CC), fluctuating light (FL), and fluctuating temperature (FT). This proof of concept evaluates whether the pipeline can resolve cultivar- and environment-associated acclimation patterns at a scale that is impractical with gas exchange alone, and whether the resulting physiological traits can identify candidates for subsequent breeding-oriented evaluation.

**Fig 1.**
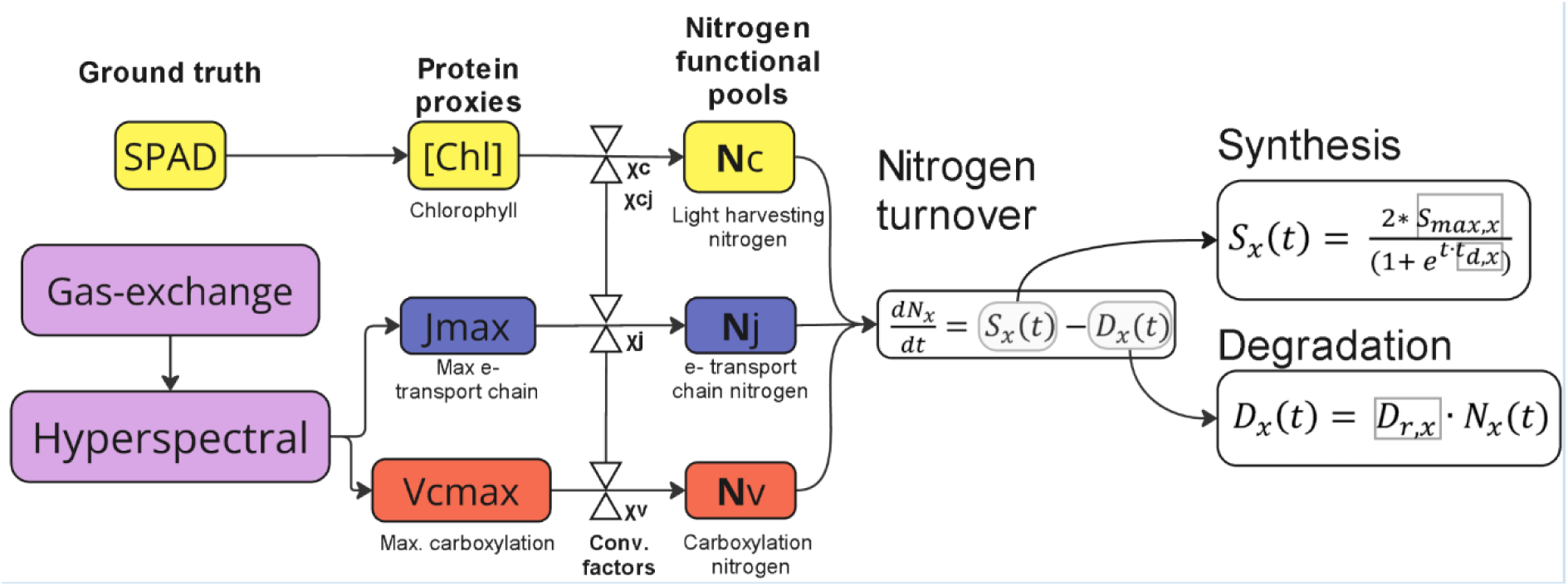
Simplified data flow of the computational pipeline proposed to high throughput phenotyping parameters related to photosynthetic acclimation, including maximal synthesis rate (*S*_max_, Equation 3), aging factor of the leaf (*t*_d,x_, Equation 3), and degradation rate (*D*_r,x_, Eq.2). Three photosynthetic nitrogen pools ([Chl]: chlorophyll concentration, *J*_max_: maximum electron transport chain, *V*_cmax_: maximum carboxylation capacity). The former was estimated from SPAD measurements, and the two latter from a hyperspectral-based model, calibrated using over two hundred *A/C*_i_ curves. Details are provided in Supplementary Figure S1.

## 2. Materials and Methods

### 2.1 Plant materials

A total of 60 winter wheat (*Triticum aestivum* L.) cultivars were selected (Supplementary Table 1): 50 representing the breeding history of winter wheat in Germany (the breeding panel) and 10 genetically diverse cultivars of exotic origin (the exotic panel). The breeding panel was derived from 191 cultivars described previously (Sabir et al., 2023; Voss-Fels et al., 2019) using a multistep selection procedure. First, five cultivars were preselected to extend or complement the available panel. Because the most recent cultivar in that dataset was released in 2013, RGT-Reform (2014), Nordkap (2016), and Asory (2018) were included to represent the subsequent decade of breeding. Two older cultivars, Impression (2005) and Julius (2008), were included because of their contrasting architectural traits. Second, only cultivars with similar heading dates in field trials were retained to reduce confounding by phenology (Lichthardt et al., 2020). Third, breeding trends were assessed by regressing grain yield under high-input field conditions against year of release using data from (Wang et al., 2025). Cultivars with a Cook’s distance greater than four times the mean were excluded to avoid overrepresentation of outliers (Cook, 1979). Genome-wide single-nucleotide polymorphism profiles were then grouped into 45 genetic clusters using k-means clustering, and the cultivar with the highest field grain yield was selected from each cluster. This procedure captured the historical range, breeding progress in grain yield, and genetic diversity of German winter wheat. The exotic panel was selected from 24 non-European cultivars using genome-wide SNP data (Lichthardt et al., 2020; Sabir et al., 2023; Wang et al., 2025). Ten cultivars representing the most genetically divergent clusters were chosen (Mabrouk et al., 2026). When a cluster contained several cultivars, selection favoured the cultivar showing the greatest plasticity in leaf mass per area, plant height, and shoot dry weight in response to planting density (Manntschke et al., 2025).

### 2.2 Growing conditions

Seeds were sown 1 cm deep in 3 × 3 × 6 cm cells (10 × 10 × 35 cm QuickPot™, Groß Kreutz, Germany) filled with bedding substrate (Substrat 1, Klasmann-Deilmann GmbH, Geeste, Germany). After 14 days of germination in a growth chamber (20 °C, 60% relative humidity, 300 µmol m⁻² s⁻¹ PPFD, and a 12-h photoperiod; Fitotron HGC 1514, Weiss Technik GmbH, Lindenstruth, Germany), the most homogeneous seedlings in which the third leaf had begun to emerge were transplanted into 10 × 10 × 35 cm QuickPot™ pots filled with the same substrate. Plants were returned to the chamber under day/night temperatures of 22/14 °C, 60% relative humidity, and a 12-h photoperiod. Incident PPFD measured 10 cm above the soil was 460 µmol m⁻² s⁻¹. Three environmental regimes were imposed: constant conditions (CC), fluctuating light (FL; hourly deviations of ±96 µmol m⁻² s⁻¹ from the mean PPFD), and fluctuating temperature (FT; hourly deviations of ±2.4 °C from the mean temperature; Fig. 2). Because the chamber could impose only one climate programme at a time, CC, FL, and FT were conducted sequentially as separate runs. All runs used the same chamber settings, photoperiod, daily light integral, and temperature sum, differing only in the imposed fluctuation pattern. Treatment was therefore confounded with run (temporal block), and differences among treatments were interpreted as associations rather than replicated chamber contrasts. The realised environment of each run (Supplementary Fig. S2) was logged continuously using the growth-chamber records for air temperature, relative humidity, and light output; three additional temperature sensors positioned 5 cm above the soil in columns 6, 12, and 18 (TinyTag, Gemini Data Loggers, UK); and two temperature, relative-humidity, and CO₂ sensors positioned in columns 1 and 24 (DKCO2-Log, Driesen & Kern, Germany). Thermal time (°Cd) was calculated as the cumulative sum of daily mean air temperature above a base temperature of 0 °C, so that all developmental comparisons are expressed on a temperature-normalised time axis.

**Fig. 2.**
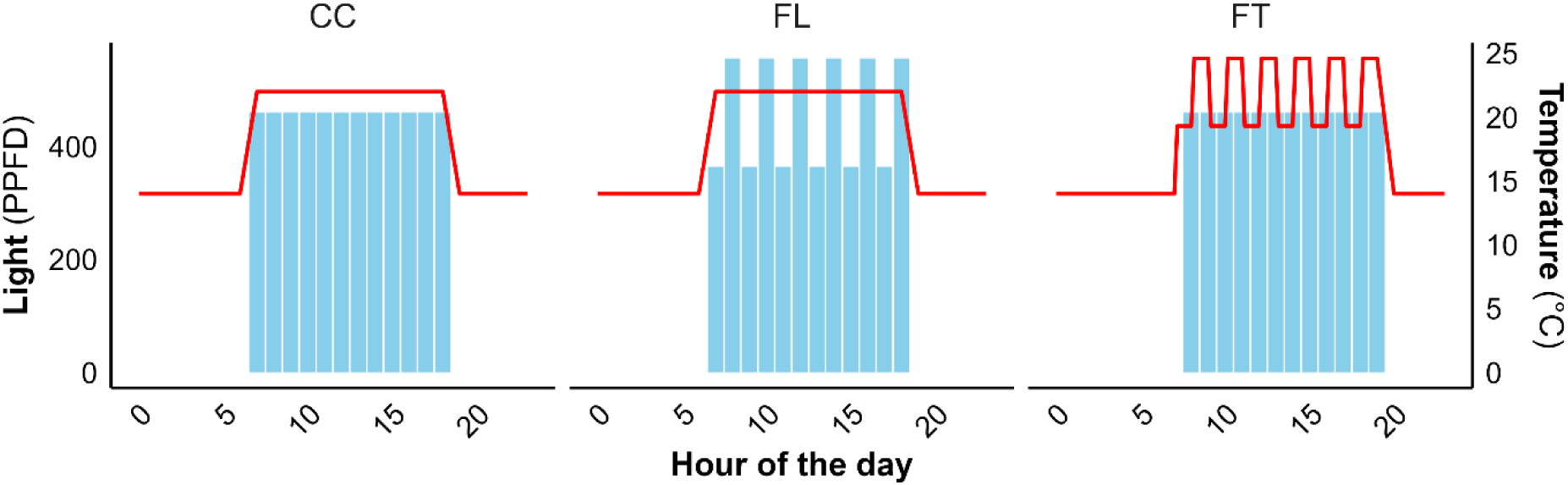
Light and temperature regimes under the three environmental conditions. Blue bars show hourly photosynthetic photon flux density (PPFD), and red lines show temperature. Treatments were constant conditions (CC; PPFD = 460 µmol m⁻² s⁻¹; day/night temperature = 22/14 °C), fluctuating light at constant temperature (FL), and fluctuating temperature at constant light (FT). In FL, PPFD varied by ±96 µmol m⁻² s⁻¹, whereas in FT, temperature varied by ±2.4 °C. Photoperiod, daily light integral, and temperature sum were identical among treatments.

During the FT run, chamber records identified a 36.1-h interruption in light between 186.9 and 215.7 °Cd. Its potential influence was assessed by simulating zero synthesis over this interval (Supplementary Fig. S2, S3).

Each treatment run followed a completely randomised block design with three spatial blocks to account for light gradients within the chamber. Randomisation was performed in *R* using its default pseudo-random number generator and a fixed seed. The three focal cultivars used for gas-exchange measurements (Durin, Diplomat, and RGT-Reform) were represented by five biological replicates distributed randomly among the blocks to maintain sampling throughout the experiment. Each of the other 57 cultivars was represented by one biological replicate per block (3 plants in total).

### 2.3 SPAD measurements

Chlorophyll content was estimated every 3–4 days from four positions along the adaxial surface of the third leaf using a SPAD-502Plus chlorophyll meter (Konica Minolta, Germany). Mean leaf chlorophyll content was calculated using wheat-specific coefficients (Parry et al., 2014). The chlorophyll content index (CCI) was first calculated from the SPAD value (*CCI* = 1 + 0.00119 · *SPAD*^2.67^) and then converted to chlorophyll content (µmol m⁻²; Chlorophyll = −84 + 79 ∗ *CCI*^0.6^). At full senescence, chlorophyll approaches zero and SPAD readings reach the instrument floor. When the device returned an error for a highly senescent measurement position, that position was assigned a value of zero before the four-position mean was calculated. This prevented the leaf mean from being biased upwards by retaining only the remaining positive readings.

### 2.4 Gas-exchange measurements and estimation of photosynthetic parameters

Gas exchange was measured across the third-leaf lifespan in three focal cultivars (Durin, Diplomat, and RGT Reform) using two LI-6800 portable photosynthesis systems (LI-COR Biosciences, Nebraska, USA). Measurements began one week after transplant and were repeated every 3–4 days until complete senescence. In total, 246 *A*/*C*_i_ curves were collected: 110 under CC on 12 dates, 68 under FL on 7 dates, and 68 under FT on 8 dates. The median number of replicate plants per cultivar × treatment × date was three (range 1–5). *A*/*C*_i_ and light-response curves were measured at the middle of the third leaf. Before each measurement, both systems were checked for leaks and potential malfunctions (Busch et al., 2024). Measurements were conducted in a growth chamber set to the same environment as the experimental chamber. Before each *A*/*C*_i_ curve, leaves were acclimated at 1300 µmol m⁻² s⁻¹ PPFD, a flow rate of 600 µmol s⁻¹, 50% relative humidity, and an air temperature of 25 °C. *A*/*C*_i_ curves were measured at twelve CO₂ concentrations (400, 300, 200, 100, 50, 150, 250, 350, 600, 900, 1200, and 1500 µmol mol⁻¹). Day respiration was estimated using the Kok method (Yin and Amthor, 2024). Light-response curves were obtained by decreasing PPFD through 200, 150, 125, 100, 75, 50, 40, 30, 20, 10, and 0 µmol m⁻² s⁻¹ at a flow rate of 300 µmol s⁻¹, 400 µmol mol⁻¹ CO₂, 50% relative humidity, and 25 °C. The transition between dark and light respiration was set at 30 µmol m⁻² s⁻¹, consistent with our observations. The resulting day respiration was used to estimate electron transport at 1300 µmol m⁻² s⁻¹ PPFD (*J*_high_) and maximum carboxylation capacity (*V*_cmax_) from the *A*/*C*_i_ curves. Two curve-fitting approaches were compared: the established *plantecophys* R package (version 1.4.6; Duursma, 2015), which uses iterative non-linear least squares, and *PhotoGEA* (Lochocki et al., 2025), which provides an analytical solution to the Farquhar–von Caemmerer–Berry model. Mesophyll conductance (*g*_m_) was not estimated because reliable chlorophyll-fluorescence measurements required for the variable-J method were unavailable for all curves. The fitted parameters therefore represent apparent values based on intercellular CO₂ concentration (*C*_i_), rather than chloroplastic rates. Default temperature-dependent values were used for the Michaelis–Menten constant (*K*_m_) and CO₂ compensation point (Γ). *J*_high_ was then adjusted following (Buckley and Diaz-Espejo, 2015) to obtain *J*_max_ (Equation 4):

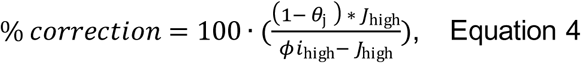

where *φi*_high_ is 0.331, representing a leaf absorptance of 0.86, from an average of 10 dicot species (Evans and Poorter, 2001) and *θ*_j_ is 0.825 (Bernacchi et al., 2003).

### 2.5 Hyperspectral measurements to estimate photosynthetic parameters

Hyperspectral reflectance was measured at the midpoint of the third leaf using a full-range spectroradiometer fitted with a leaf clip and dark background (ASD FieldSpec 4, Malvern Panalytical Ltd, UK). For the focal gas-exchange measurements, spectra were collected on the same day, with no more than 3 h between measurements. Across the experiment, reflectance from 350 to 2500 nm was measured twice per week for 5 weeks after leaf appearance. The instrument had a spectral resolution of 3 nm in the visible and near-infrared region (350–1000 nm) and 8 nm in the shortwave-infrared region (1000–2500 nm), with sensor transitions at 1000 and 1800 nm. White references were collected using Spectralon at least every 5 min. Each record was averaged from at least five 100-ms scans; additional scans were taken when needed until reflectance no longer shifted visibly. The instrument was allowed to acclimate to the surrounding temperature for at least 25 min before measurement (Burnett et al., 2021). Reflectance spectra were smoothed and corrected for sensor-transition discontinuities using the *hsdar* package in R (Lehnert et al., 2019).

A dataset of 245 matched gas-exchange and hyperspectral observations from the three focal cultivars was used to train partial least squares regression (PLSR) models with the pls package (version 2.8.3; Mevik and Wehrens, 2007). The models predicted *V*_cmax_ and *J*_high_, with *J*_high_ subsequently converted to *J*_max_ using Equation 4, for all experimental plants, including those without direct gas exchange, following the practices described by Burnett et al. (2021) and Fu et al.(2019). Six spectral ranges were evaluated: 350–2500, 400–2150, 500–2000, 400–1000, 700–2150, and 1300–2400 nm (Supplementary Fig. S4). Up to 20 PLSR components were tested to limit overfitting. Internal validation used stratified 5-fold cross-validation, with four folds for calibration and one for validation (an 80/20 split). Stratification balanced genotype and approximately 200 °Cd life-stage intervals across folds. Each individual observation was assigned to only one-fold and therefore did not occur in both calibration and validation. However, because the same plants were measured repeatedly, observations from different dates on the same plant could occur in both subsets. Results from the five validation folds were combined (Fig. S5), and stability was assessed across five random seeds.

### 2.6 Conversion of photosynthetic capacity proxies to nitrogen pools

Nitrogen invested in carboxylation (*N*_v_, mmol N m⁻²; Equation 5), electron transport (*N*_j_, mmol N m⁻²; Equation 6), and light harvesting (*N*_c_, mmol N m⁻²; Equation 7) was estimated following (Buckley et al., 2013):

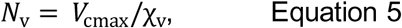

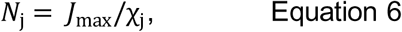

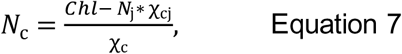

where χ*_v_* (4.49 μmol CO_2_ mmol^−1^ N s^−1^) is the carboxylation capacity per unit Rubisco nitrogen, χ*_j_* (9.48 μmol e^-^ mmol^−1^ N s^−1^) is the electron transport capacity per unit electron transport nitrogen, and χ*_cj_* (4.64 × 10⁻⁴ mmol Chl mmol^-1^ N) and χ*_c_* (0.034 mmol Chl mmol^-1^ N) are the conversion coefficients for chlorophyll per electron transport nitrogen and per light-harvesting component nitrogen, respectively. Since hyperspectral and chlorophyll measurements were sometimes taken on consecutive days, *N*_v_ and *N*_j_ (derived from PLSR-predicted *V*_cmax_ and *J*_max_) were linearly interpolated to match the thermal time of each SPAD chlorophyll measurement. This temporal alignment enabled calculation of light-harvesting nitrogen (*N*_c_) using the partitioning equation at each chlorophyll measurement point.

Photosynthetic nitrogen (*N*_ph_, mmol N m⁻²) was defined as biologically active nitrogen invested in proteins associated with carboxylation, electron transport, and light harvesting. It was calculated as the sum of the three pools (Equation 8):

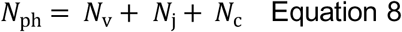

The photosynthetic nitrogen partitioning fraction of pool x (*p*_x_) was calculated as the ratio of nitrogen in that pool (*N*_x_, mmol N m⁻²) to *N*_ph_:

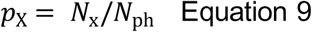

### 2.7 Parameterization of the turnover model

Chlorophyll and nitrogen-pool dynamics were parameterised using the turnover model described in Equations 1–3 over thermal time (°Cd). To compare chlorophyll and nitrogen-pool parameters without changing the model structure, chlorophyll concentrations were rescaled by a factor of 10 to match the approximate magnitude of the nitrogen pools. The model was fitted separately for each cultivar × environment combination using the *DEoptim* global optimisation algorithm (Mullen et al., 2011), minimising the root-mean-square deviation between estimated and simulated concentrations (mmol m⁻²). Simulations used the LSODA algorithm implemented in the *deSolve* R package (Soetaert et al., 2010), with a time step of 0.4 °Cd. Parameter bounds followed previous work (Pao et al., 2020). To improve the fit and ensure biologically realistic boundary conditions, artificial zero values were introduced at thermal times 0 and 700 °Cd. These points reflect known physiological states where the parameter value is zero despite the absence of direct measurements at these time points. A two-stage optimisation procedure was used, following Pao et al.(2020, 2019a), to reduce convergence on local minima and limit parameter trade-offs in sparsely sampled trajectories. In Stage 1, *S*_max_, *t*_d_, and *D*_r_ were fitted freely over broad parameter ranges. In Stage 1, all three parameters (*S*_max_, *t*_d_, *D*_r_) were allowed to vary in a broad parameter search to assess whether timing parameters (*t*_d_ and *D*_r_) exhibited meaningful variation rather than assuming genotype-invariant values, as originally done by Pao et al. (2020). In Stage 2, *t*_d_ and *D*_r_ were fixed at their Stage 1 values, and optimization focused on *S*_max_, the parameter of primary biological interest and expected genotypic variability. The *DEoptim* algorithm’s population-based approach inherently explores multiple starting points within the parameter space, reducing sensitivity to initial conditions.

### 2.8 Uncertainty assessment and parameter identifiability

Because uncertainty enters at several stages of the pipeline, we evaluated five sources of uncertainty and errors (ERR-1 to ERR-5) using complementary analyses (Supplementary Fig. S1). ERR-1 evaluates the effect of the fixed biochemical conversion coefficients (*χ*_v_, *χ*_j_, *χ*_c_ and *χ*_cj_, Equation 5-7), including the dependence of *N*_c_ on *N_j_*. It was assessed by coefficient sensitivity across all 60 cultivars. Each of the four coefficients was varied separately at seven levels spanning a conservative ±25% envelope, giving 28 perturbed coefficient sets. After each perturbation the nitrogen pools were recalculated, the turnover model refitted, and the resulting standard deviations in *S*_max_, *t*_d_ and *D*_r_ were recorded. Conclusion stability was then assessed by recalculating the principal biological results for every perturbed set and recording whether their original sign and significance were preserved, comprising 1,764 checks: 756 release-year associations, 756 cross-environment correlations and 252 treatment rankings.

ERR-2 evaluates the prediction error of the PLSR models for *V*_cmax_ and *J*_high_. It was quantified using the measurement-level validation described in Section 2.5, repeated across five random seeds, and summarised as the held-out R^2^ and RMSE of each model (Supplementary Fig. S5). Because the same plants were measured repeatedly, observations from different dates on the same plant could occur in both the calibration and validation subsets. ERR-2 therefore quantifies measurement-level prediction error and is not independent at the plant or cultivar level, which motivates ERR-3.

ERR-3 evaluates the potential effects of extrapolation when the PLSR models are transferred from the three calibration cultivars to the full 60-cultivar panel. It was assessed by leave-one-genotype-out (LOGO) cross-validation, in which each calibration cultivar was held out in turn and the models were retrained on the remaining two before predicting the held-out cultivar (Supplementary Figs. S6, S7). The resulting LOGO error was adopted as the conservative between-cultivar error bound, and used to set the prediction-screening tolerances (Section 3.3). Two additional checks evaluated the transfer: i) a cultivar learning curve, in which held-out RMSE was compared between calibration sets of one and two cultivars (Supplementary Fig. S8) and ii) a comparison of spectral space, in which principal component analysis of leaf reflectance tested whether observations from the remaining cultivars fell within the convex hull of the calibration cultivars in PC1–PC2 space (Supplementary Fig. S9). Residual behaviour was examined against predicted value, treatment and leaf age (Supplementary Fig. S10).

ERR-4 evaluates the error in fitting the turnover model to the pool trajectories. It was assessed by residual bootstrapping in 540 conditions comprising all 60 cultivars × three treatments × three pools. Within each trajectory, raw residuals (observed minus predicted) from real observations were sampled independently with replacement, without centring, and added to the central fitted values. Each trajectory was refitted 100 times (54,000 refits in total) by bounded nonlinear least squares using the Levenberg–Marquardt algorithm, initialised at the original parameter estimates. Bounds wider than those of the original fits were used so that the bootstrap draws could not pile up against the edge of the search box. For *S*_max_, *t*_d_ and *D*_r_, respectively, lower bounds of 0, −0.1 and 0, and upper bounds of 6 × *S*_max_ + 1 (minimum 5), 0.1 and 1. Parameter distributions were summarised as percentile 95% intervals (2.5th and 97.5th percentiles) and coefficients of variation for each pool and treatment (Supplementary Fig. S14).

ERR-5 evaluates the identifiability and recovery of *S*_max_, *t*_d_ and *D*_r_. It was assessed by synthetic-data recovery. For every cultivar × treatment × pool combination, the fitted parameter values were treated as true values and used to simulate trajectories at the actual measurement times in the emergence frame. Gaussian noise was added at the residual SD of the corresponding pool, using one noise level per pool estimated from the residuals of all fits for that pool (Supplementary Fig. S15); the synthetic zero points used in the original fitting were excluded. Each synthetic trajectory was refitted by bounded Levenberg–Marquardt minimisation warm-started at the true values, using the bounds of the original fits (*S*_max_: 0.01–2; *t*_d_: 0.0001–0.015; *D*_r_: 0.001–0.03), with 15 replicate refits per condition (8,100 refits in total). The differential-evolution stage of the two-stage procedure was not repeated; this analysis therefore assessed local identifiability under observational noise rather than recovery from arbitrary starting values. A recovered estimate was classified as reaching a bound when it equalled that bound to within a relative distance of 10⁻⁶ (Supplementary Fig. S15).

We estimated how much two additional inputs could vary but did not test how this variation affected the full analysis. They are therefore reported as limitations rather than quantified uncertainty terms: the SPAD-to-chlorophyll conversion (Gaussian, SD = 49 µmol m⁻²; wheat calibration of Parry et al. (2014), R² = 0.87) and the leaf-age zero point (Gaussian, SD = 8.5 °Cd). These two terms were not propagated separately. The random part of the SPAD error is already contained in the residuals used for ERR-4, and a systematic error in the SPAD calibration enters Nc in approximately the same way as the χc coefficient varied in ERR -1. The leaf-age zero point remains untested, but the same offset is applied to all cultivars within a treatment, so it can shift absolute timing without affecting comparisons among cultivars.

### 2.9 Data analysis

Leaf age was expressed relative to third-leaf emergence. Because emergence could not be recorded directly for every plant, it was estimated separately for each treatment from chlorophyll accumulation. For each genotype × treatment combination, observations up to peak chlorophyll (the rising limb) were fitted with a synthesis-only form of the turnover model. The onset parameter *t*_0_ was fitted freely, while degradation was disabled because it was assumed to be negligible during the accumulation phase. Differential-evolution optimisation constrained *t*_0_ to occur at or before the first measurement. The median *t*_0_ within each treatment was used as the emergence offset and subtracted from thermal-time leaf age. The same offset was applied to chlorophyll, the three nitrogen pools, and gas-exchange capacities so that observations and fitted curves shared a common time axis from emergence.

All analyses and visualisations were performed in R version 4.3.1. Associations between treatment and turnover parameters were assessed by one-way ANOVA followed by Tukey’s HSD test (*p* < 0.05). Pearson correlations were used to assess (i) associations between parameter values and cultivar release year and (ii) cross-environment stability by relating parameter values for the same cultivar across treatments. Release-year analyses included all 50 cultivars in the German breeding panel; exotic accessions were excluded. Reported p values were not adjusted for multiple testing and were interpreted as exploratory. Correlations were calculated and interpreted separately for *N*_c_, *N*_j_, and *N*_v_ because the pools are not independent: *N*_c_ is derived from *N*_j_, while *N*_j_ and *N*_v_ share the same gas-exchange fit.

## 3. Results

### 3.1 A computational pipeline integrating rapid optical sensing with a mechanistic model of photosynthetic protein-turnover allows robust high-throughput characterization of photosynthetic acclimation

Two algorithms, *PhotoGEA* and *plantecophys*, were used to obtain *V*_cmax_ and *J*_high_ from gas-exchange data. *PhotoGEA* avoids convergence problems that may occur with noisy or atypical curves, particularly in senescing leaves (Lochocki et al., 2025). It returned *V*_cmax_ for more curves than *plantecophys* (n = 241 versus 195). Its lower *J*_high_ count (n = 171) reflects reporting logic rather than fit quality: *PhotoGEA* reports *J*_high_ only when a curve contains electron-transport-limited points, whereas *plantecophys* reports both *V*_cmax_ and *J*_high_ for every fitted curve (n = 195). To maintain biological consistency when nitrogen pools were calculated, *V*_cmax_ and *J*_high_ predictions were required to use the same gas-exchange fitting method (*PhotoGEA* or *plantecophys*). Models were selected using the mean of the cross-validated R² values for both parameters. *PhotoGEA* gave the best combined performance (mean R²: *plantecophys* = 0.483; *PhotoGEA* = 0.528). The optimal spectral range was 500–2000 nm for *V*_cmax_ (R² = 0.56 ± 0.14 across seeds; RMSE = 27.0 µmol CO₂ m⁻² s⁻¹; n = 241) and 700–2150 nm for *J*_high_ (R² = 0.50 ± 0.18; RMSE = 38.3 µmol m⁻² s⁻¹; n = 171).

### 3.2 Chlorophyll dynamics of leaves show cultivar-specific responses to environmental fluctuations

After leaf appearance, chlorophyll content increased during early development (leaf expansion), reached a peak around 150–250 °Cd, and subsequently declined during senescence (Fig. 3A). This general trajectory was consistent across genotypes, reflecting the balance between chlorophyll synthesis during expansion and degradation during senescence. Clear differences in curve shape nonetheless emerged among environments. Leaves under fluctuating light (FL) reached the highest peak chlorophyll content, leaves under constant conditions (CC) senesced earliest and most steeply, and those under fluctuating temperature (FT) retained chlorophyll the longest (Fig. 3A). Fitting these curves per genotype and environment (Eqs. 1–3) resolved this ordering into three turnover parameters, which differed significantly among all three environments (Tukey HSD, *p* < 0.05). CC showed the highest maximum synthesis rate (*S*_max_; Fig. 3B), the fastest age-related decline in synthesis (highest *t*_d_; Fig. 3C), and the highest degradation rate (*D*_r_; Fig. 3D). FT showed the lowest *S*_max_, *t*_d_, and *D*_r_ (Fig. 3B–D), so its slower degradation and more sustained synthesis extended greenness despite slower chlorophyll accumulation. FL was intermediate in all three parameters. The FT rising limb also showed a transient interruption near 186.9 °Cd (Fig. 3A), coinciding with a measured ∼36 h light interruption in the growth chamber. A forward simulation that disabled synthesis for the measured duration slightly reduced RMSD (11.3 versus 12.9; Supplementary Fig. S3).

**Fig. 3.**
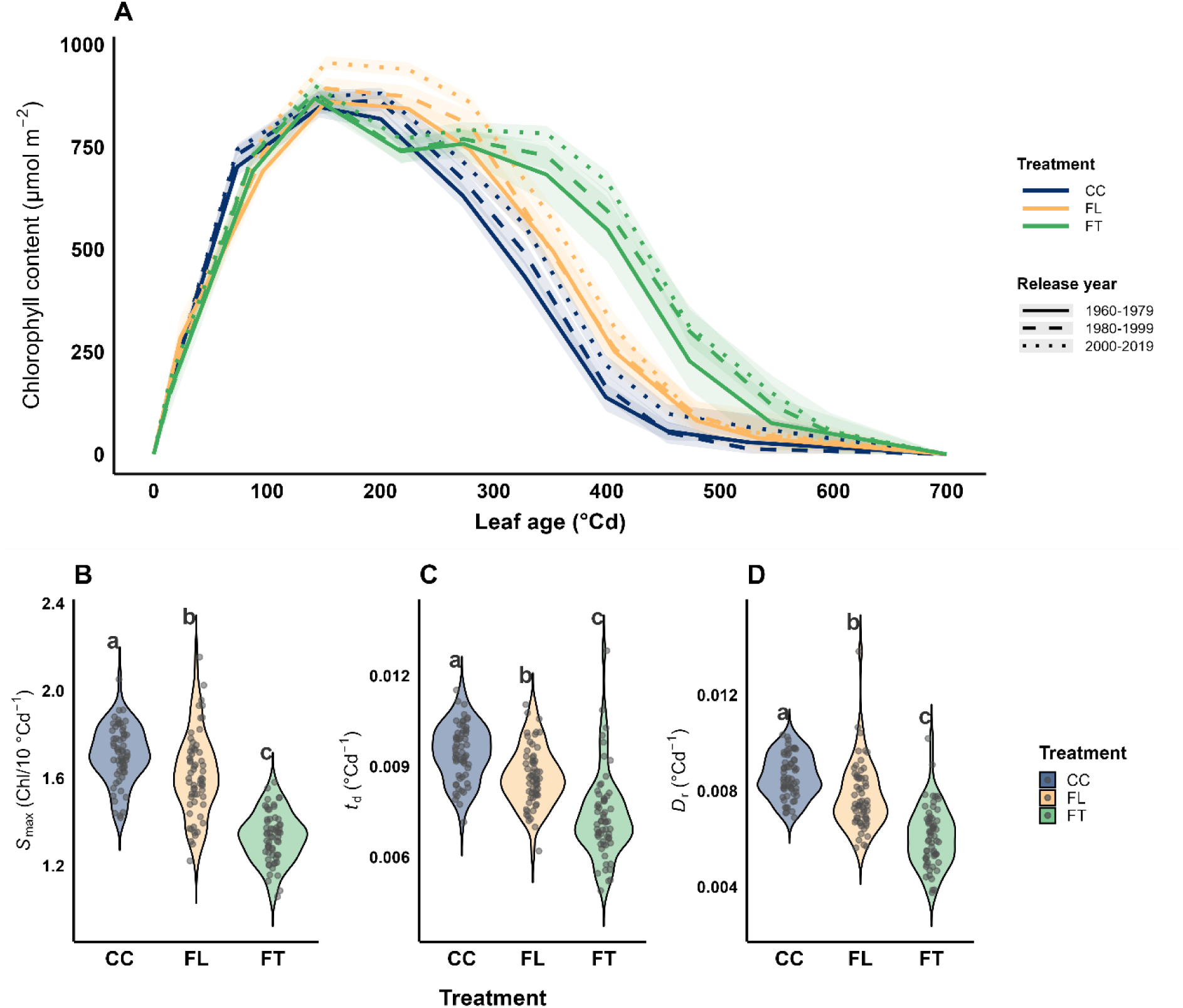
Dynamics of chlorophyll concentration of the third leaf (A) and parameters of chlorophyll turnover (B-D) under constant light and temperature (CC), fluctuating light and constant temperature (FL) and constant light and fluctuating temperature (FT, Fig. 1). Line pattern indicates the era in which the cultivar was released, the shadow represents the standard deviation, and the colour indicates the different treatments. Parameters include (B) maximum chlorophyll synthesis (*S*_max_, Equation 3), (C) aging factor (*t*_d_, Equation 3), and (D) degradation rate (*D*_r_, Equation 2). Letters indicate significant differences between treatments (Tukey HSD, *p* < 0.05). CC shows the highest values between treatments in all parameters. Dots in (B)-(D) represent individual cultivars; the violin shows the distribution across all 60 cultivars.

### 3.3 Hyperspectral reflectance enables high-throughput phenotyping of dynamic photosynthetic protein turnover

Hyperspectral predictions of *V*_cmax_ *and J*_high_ using PLSR showed moderate accuracy (ERR-2; *R²* = 0.56 and 0.50, respectively) with minimal bias (−1.44 and −2.44; Supplementary Fig. S5). Prediction errors were randomly distributed and showed no significant systematic dependence on leaf age, predicted value, cultivar, or environmental conditions (Fig. S10), enabling phenotyping across all cultivars and time points.

Before extrapolating the PLSR models calibrated on three cultivars with direct gas-exchange measurements to the remaining 57 cultivars, we first assessed whether their spectral properties fell within the calibration domain (ERR-3, Supplementary Fig. S1). Ninety-nine percent of spectra from the 57 additional cultivars fell within the calibration convex hull in PC1–PC2 space (Supplementary Fig. S9), providing strong evidence that the models were applied within their spectral domain. LOGO cross-validation further supported the robustness of the extrapolation, yielding *R²* = 0.44 for *V*_cmax_ and 0.43 for *J*_high_, with no pronounced cultivar-specific offsets (Supplementary Figs. S6, S7, S8). Moreover, expanding the calibration set from one to two cultivars reduced RMSE, particularly for *J*_high_, indicating that the models captured transferable spectral–physiological relationships rather than cultivar-specific patterns. Together, these complementary tests provide strong support for extending the PLSR models to the full set of genotypes in the experiment.

Outlier detection was applied independently to *V*_cmax_ and *J*_high_. First, values below a one-sided tolerance of 1.5×LOGO RMSE/2 (*V*_cmax_ <−22.9 µmol CO₂ m⁻² s⁻¹; *J*_high_ <− 36.1 µmol e⁻ m⁻² s⁻¹) or values above biologically plausible upper bounds (*V*_cmax_ > 300; *J*_max_ > 450 after conversion using Equation 4), were excluded. This excluded 1.4% and 3.8% of predictions, respectively, mostly during late senescence. Second, within-treatment LOESS curves were fitted (span = 0.75) and observations were removed when their residuals exceeded either three treatment-specific standard deviations or 1.5 times the LOGO RMSE. This excluded a further 4.5% and 7.0%, respectively, across leaf development (Supplementary Fig. S11).

Observations were classified as outliers by either of two rules: a residual exceeding three standard deviations of the per-treatment residual distribution, or a residual exceeding an absolute cap derived as 1.5 × the leave-one-genotype-out RMSE. Across the panel, the prediction-tolerance filter removed 1.4% of *V*_cmax_ and 3.8% of *J*_high_ predictions, almost entirely from the senescent tail where capacity approached zero. The LOESS procedure removed a further 4.5% and 7.0%, respectively, distributed across leaf development (Supplementary Fig. S11).Predictions below these tolerances, or values above biologically plausible upper bounds (*V*_cmax_ > 300; *J*_max_ > 450 after conversion using Equation 4), were excluded.

In direct gas-exchange measurements, *V*_cmax_ and *J*_max_ increased rapidly after emergence and declined with leaf age under all treatments (Fig. 4A, C). CC reached earlier peaks, whereas FT retained higher photosynthetic capacity later in the leaf lifespan. The hyperspectral estimates reproduced this temporal structure across 60 cultivars (Fig. 4B, D), with peaks around 150–250 °Cd followed by a senescence-related decline. FT generally retained *J*_max_ and *V*_cmax_ later, while FL showed lower *J*_max_ through much of the lifespan.

**Fig. 4.**
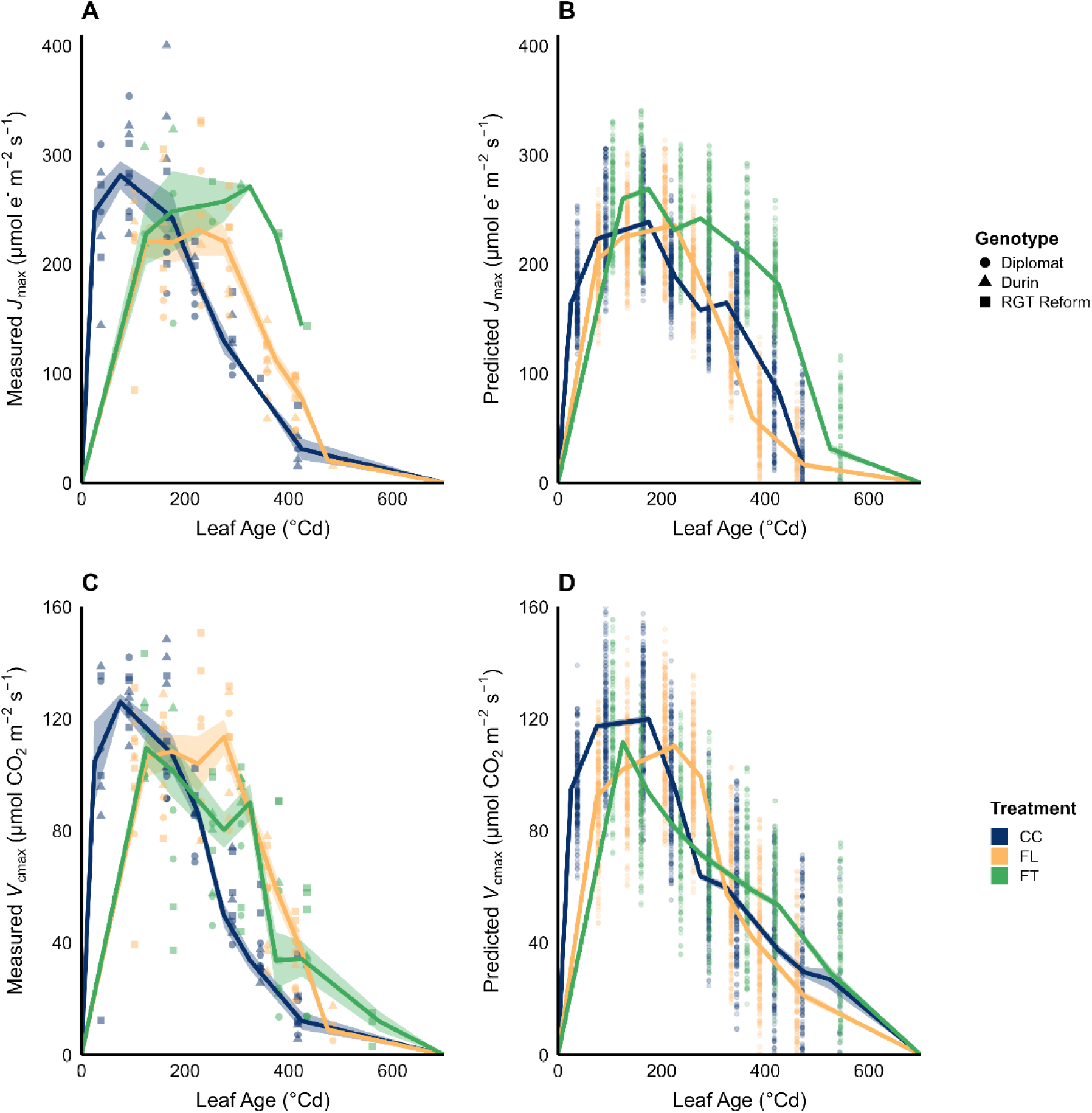
Measured and hyperspectral-predicted photosynthetic capacity across leaf age. (A, C) *J*_max_ and *V*_cmax_ measured by gas exchange in three focal cultivars (Diplomat, Durin, and RGT Reform); symbols identify cultivars. (B, D) Corresponding hyperspectral predictions across 60 cultivars. Colours indicate constant conditions (CC), fluctuating light (FL), and fluctuating temperature (FT). Lines summarise treatment-specific trajectories, and shaded bands in the measured panels show variability around the trajectory. Each point represents one measurement. Synthetic zero anchors at 0 and 700 °Cd, used to constrain turnover-model fitting, are not direct measurements; outliers were removed using the prediction-tolerance and LOESS rules described in Methods.

Using the estimated chlorophyll content (Fig. 3A), *J*_max_ (Fig. 4B), and *V*_cmax_ (Fig. 4D), we converted them to leaf nitrogen content by published coefficients (Eqn. 5-7, Supplementary Table 2) and estimated the dynamic of total nitrogen content per leaf area and dynamic photosynthetic nitrogen partition in light-harvesting (*N*_c_), electron transport (*N*_j_), and carboxylation (*N*_v_) pools (Fig. 5).

**Fig 5.**
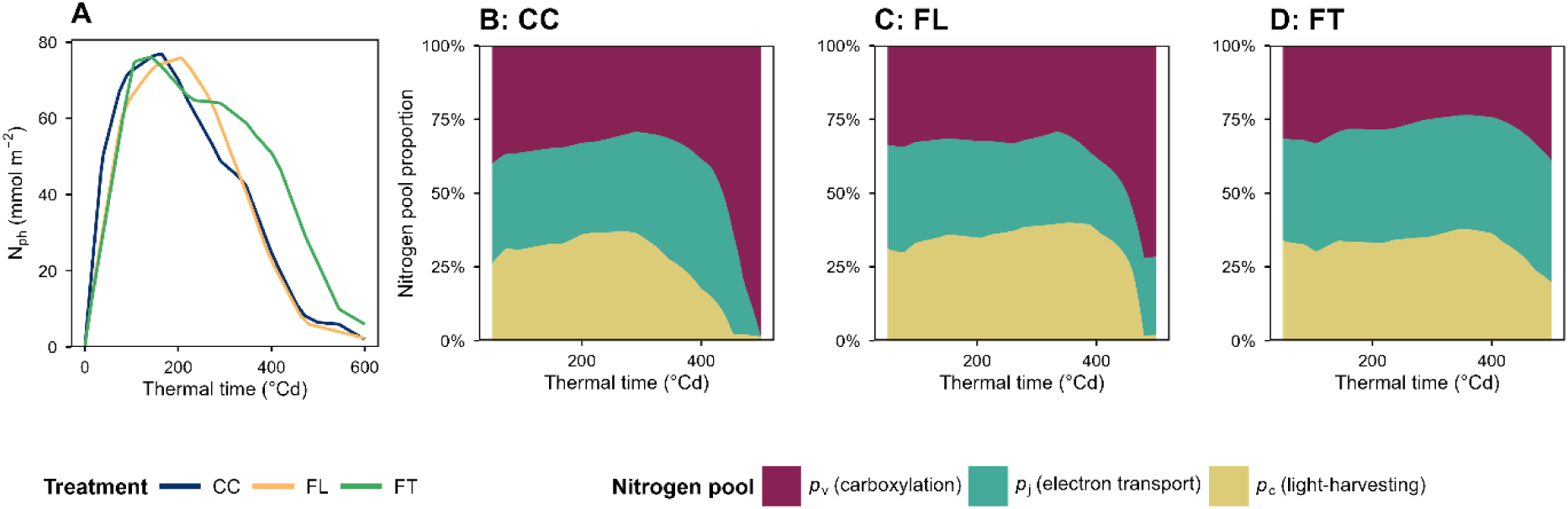
A. Mean total photosynthetic nitrogen of 60 winter wheat cultivars over time for each environmental condition. **B-D.** Proportional contributions to *N*_ph_ of three nitrogen pools, *p*_c_ (light-harvesting N pool), *p*_j_ (electron transport N pool), and *p*_v_ (carboxylation N pool), to *N*_ph_ across thermal time (°Cd) for (**B**) constant temperature and light (CC), (**C**) fluctuating light (panel C, FL), and (**D**) fluctuating temperature (FT). Panels B-D cover thermal time 60-560 **°**Cd to avoid artifacts in the ratio calculation when the total nitrogen is small.

Total photosynthetic nitrogen (*N*_ph_) followed a similar developmental trajectory to chlorophyll content, peaking at ∼70 mmol m⁻² around 150-250 °Cd before declining during senescence (Fig. 5A). Total nitrogen remained highest in the FT run throughout most of the second half of leaf lifespan, CC had the fastest decrease in *N*_ph_, and FL the slowest increase. The proportional contributions of the three nitrogen pools changed systematically with leaf age (Fig. 5B–D). During early leaf development (60–200 °Cd), the fraction allocated to light harvesting (*p*_c_, Eq. 9) was initially the smallest but gradually increased to approximately 30% of total *N*_ph_ across all treatments. This increase occurred at the expense of the electron transport (*p*_j_) and carboxylation (*p*_v_) pools. This trend continued until approximately 300 °Cd under CC and 400 °Cd under both FL and FT, after which *p*_c_ was preferentially degraded. Under FL, the decline was most rapid. Under FT, *p*_c_ decreased more gradually and remained above 20% of total *N*_ph_ until nearly 500 °Cd, whereas under CC and FL it fell to almost zero between 450 and 500 °Cd. The electron transport pool (*p*_j_) followed a similar trajectory but consistently represented a larger fraction of *N*_ph_, peaking at 35–45% around 250–350 °Cd. Its decline lagged behind that of *p*_c_, making *p*_j_ the last non-carboxylation pool to be degraded. As with *p*_c_, degradation of *p*_j_ was slower under FT than under CC or FL. As *p*_c_ and *p*_j_ declined, the carboxylation pool (*p*_v_) increased correspondingly. Under CC and FL, *p*_v_ approached 100% of total *N*_ph_ by approximately 500 °Cd, whereas under FT *p*_c,_ and especially *p*_j,_ persisted beyond 500 °Cd.

### 3.4 Turnover-model parameters reveal pool-specific acclimation patterns across environmental treatments

The estimated dynamics of the light-harvesting (*N*_c_), electron-transport (*N*_j_), and carboxylation (*N*_v_) nitrogen pools were fitted with the mechanistic turnover model (Eqs. 1–3). For each cultivar, the model returned three biologically interpretable parameters: maximum synthesis rate (*S*_max_), age-related decline in synthesis (*t*_d_), and degradation rate (*D*_r_; Fig. 6). Sensitivity analysis (ERR-1; Supplementary Figs. S1, S12) showed that uncertainty in the biochemical conversion coefficients (Eqs 5-8) mainly affected the magnitude of *S*_max_ (relative SD up to ∼19%), while effects on *t*_d_ and *D*_r_ were <1%. Importantly, none of our biological conclusions changed when the conversion coefficients were varied. Across the 28 parameters perturbations, all biological claims (treatment comparisons, cross-environment and release year correlations) gave the same answer as with the published coefficients, in both direction and statistical significance. Comparison of predicted photosynthetic protein turnover with PLSR estimates showed residuals centred around zero (Supplementary Fig. S13), with a slight temporal pattern of overestimation early and late development of the leaves and underestimation in the middle. Despite these slight deviations, the overall trajectory and treatment patterns were preserved, indicating that the model captures the biologically relevant differences among treatments and developmental stages. Before comparing genotype and treatment effects on parameters of photosynthetic protein turnover, we evaluated their precision and identifiability (ERR-4 and ERR-5, Supplementary Fig. S1). Across the full panel, residual bootstrapping yielded median coefficients of variation of 10.3%, 14.8%, and 19.6% for *S*_max_; 20.9%, 24.6%, and 26.0% for *t*_d_; and 25.9%, 32.5%, and 34.9% for *D*_r_ across *N*_c_, *N*_j_, and *N*_v_, respectively. All 54,000 refits converged, with precision highest for *S*_max_ and lowest for *D*_r_, particularly for *N*_v_ (Supplementary Fig. S14). Synthetic-data recovery confirmed these patterns: fitted relationships were generally close to 1:1 and unbiased to slightly positive, with 63%, 41%, and 38% of *S*_max_, *t*_d_, and *D*_r_ estimates, respectively, within ±20% of the true value. *S*_max_ never reached an optimisation bound, whereas *t*_d_, and *D*_r_ did so in 11.4% and 9.1% of refits, respectively, most frequently for *N*_v_ under FT (Supplementary Fig. S15). Together with the positive *S*_max_–*D*_r_ and negative *t*_d_–*D*_r_ correlations (Supplementary Fig. S16), these results identify *D*_r_ as the least precisely estimated parameter, but also show that the model is sufficiently identifiable to support robust comparative inference, even where absolute parameter estimates warrant caution.

**Fig. 6.**
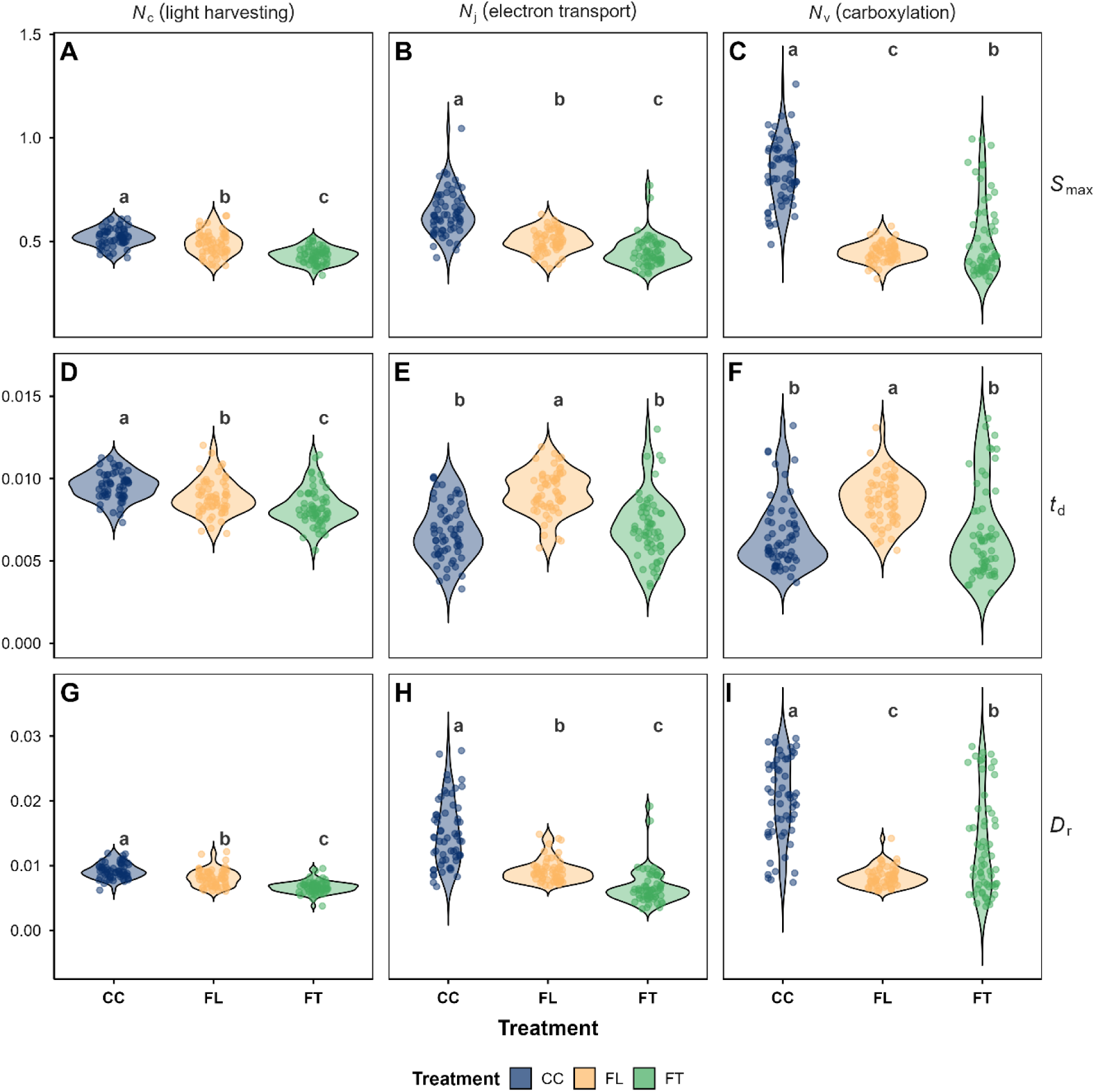
Treatment-associated variation in turnover-model parameters for the light-harvesting (*N*_c_; A, D, G), electron-transport (*N*_j_; B, E, H), and carboxylation (*N*_v_ ; C, F, I) nitrogen pools. Rows show maximum synthesis rate (*S*_max_; A–C), age-related decline in synthesis (*t*_d_; D–F), and degradation rate (*D*_r_; G–I). Points represent individual cultivars, and violins show the distributions across 60 cultivars under constant conditions (CC), fluctuating light (FL), and fluctuating temperature (FT). Different letters indicate significant differences among treatments within each panel (Tukey HSD, *p* < 0.05).

**Fig. 7.**
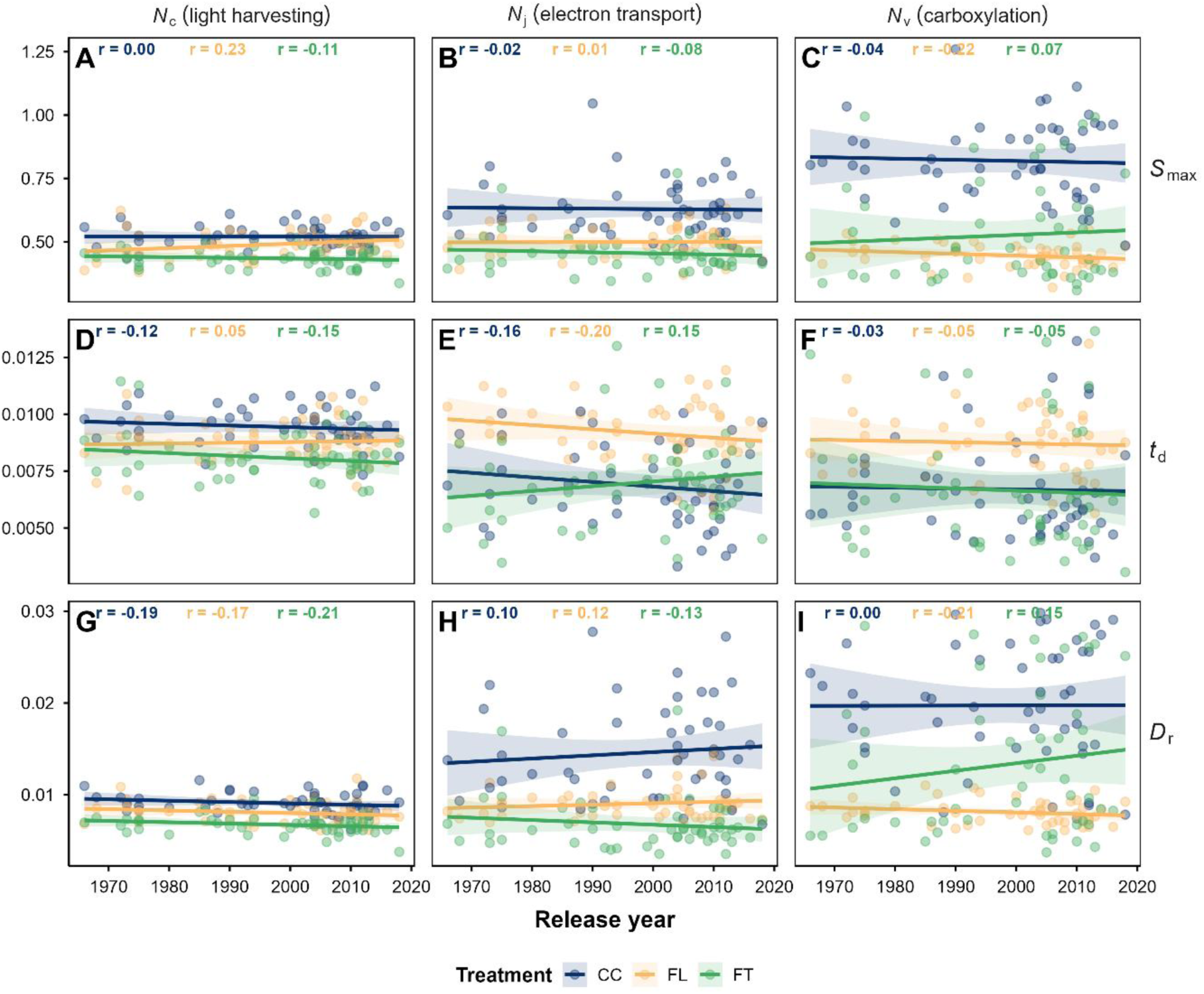
Associations between cultivar release year and turnover-model parameters for the light-harvesting (*N*_c_; A, D, G), electron-transport (*N*_j_; B, E, H), and carboxylation (*N*_v_ ; C, F, I) nitrogen pools. Rows show maximum synthesis rate (*S*_max_; A–C), age-related decline in synthesis (*t*_d_; D–F), and degradation rate (*D*_r_; G–I). Points represent the 50 cultivars with complete parameter estimates and a recorded release year in the breeding panel, excluding exotic accessions; colours indicate constant conditions (CC), fluctuating light (FL), and fluctuating temperature (FT). Lines show treatment -specific linear regressions with 95% confidence intervals, and Pearson *r* is reported at the top of each panel. None of the plotted correlations was significant at raw *p* < 0.05.

Light and temperature regimes affected the parameters of photosynthetic turnover (Fig. 6). For *N*_c_, as observed for chlorophyll dynamics in Fig. 3, *S*_max_, *t*_d_ and *D*_r_ decreased from CC to FL and then to FT (Fig. 6A, D, G). For *N*_j_, *S*_max_ and *D*_r_ also decreased from CC to FL and FT, while *t*_d_ was higher under FL than under CC and FT (Fig. 6B, E, H). For *N*_v_, *S*_max_ and *D*_r_ were highest under CC, intermediate under FT, and lowest under FL, whereas *t*_d_ was higher under FL than under CC and FT (Fig. 6C, F, I). *N*_v_ also showed substantial variation among cultivars, particularly under CC and FT. Given the single-chamber sequential design, these patterns are interpreted as treatment-level associations rather than causal effects of the fluctuation regimes.

Within each pool and treatment, *S*_max_ was positively correlated with *D*_r_, whereas *t*_d_ was negatively correlated with *D*_r_, with the strongest dependencies observed for *N*_j_ and *N*_v_ (Supplementary Fig. S16). Cross-environment correlations of the turnover parameters were weak, and their strength differed among pools (Supplementary Figs. S17–S19). Across the 27 pool × parameter × treatment-pair combinations, six were significant, all positive and all confined to the light-harvesting and electron-transport pools: *N*_c_ *S*_max_ between CC and FL (r = 0.35, raw p = 0.006) and between CC and FT (r = 0.29, raw p = 0.025), *N*_c_ *t*_d_ between CC and FT (r = 0.33, raw p = 0.009), *N*_c_ *D*_r_ between CC and FL (r = 0.31, raw p = 0.017), and *N*_j_ *t*_d_ (r = 0.32, raw p = 0.014) and *N*_j_ *D*_r_ (r = 0.28, raw p = 0.033) between CC and FL. No comparison for the carboxylation pool (*N*_v_) was significant (all |r| < 0.17). Even the strongest of these correlations explained only 12% of the between-cultivar variance, so cultivar rankings were only partly conserved between environments. The turnover parameters therefore appear largely environment-specific rather than intrinsic cultivar traits, with light-harvesting and electron-transport dynamics the partial exceptions.

To test whether photosynthetic acclimation changed across breeding history, we related *S*_max_, *t*_d_, and *D*_r_ to cultivar release year for the 50 cultivars in the German breeding panel with complete parameter estimates (Fig. 7). Pearson correlations were calculated for every nitrogen pool, parameter, and treatment. Across the 27 pool × parameter × treatment combinations, correlations with release year were weak (|*r*| ≤ 0.23) and all were non-significant (n = 50, all raw p > 0.10). The strongest tendencies were observed for *S*_max_ under FL (*N*_c_: r = 0.23, p = 0.11; *N*_v_ : r = −0.22, p = 0.12) and *D*_r_ under fluctuating conditions (*N*_c_ under FT: r = −0.21, p = 0.14; *N*_v_ under FL: r = −0.21, p = 0.15). The inferred nitrogen-pool parameters therefore provide no evidence of a directional change with cultivar release year in this panel. When treatments were pooled, parameter dispersion also did not narrow systematically across release decades (Supplementary Fig. S20), indicating no clear convergence of acclimation traits in recent cultivars.

## 4. Discussion

Photosynthesis is the fundamental physiological process sustaining plant growth and food production. Traditional gas-exchange measurements are slow (5–60 minutes per measurement), typically including 5–20 minutes for acclimation to high light, 30–40 minutes for CO2-response curves to estimate *J*_max_ and *V*_cmax_, and at least 20 minutes for a light response curve for estimating quantum yield and respiration rate (Busch et al., 2024). Furthermore, these measurements cover only a small area of a single leaf, making it particularly time-consuming to capture photosynthetic acclimation across environmental conditions and leaf ageing because repeated measurements are needed (Calzadilla et al., 2022). The main advance of this study is the capacity to phenotype photosynthetic acclimation at scale using rapid and comparatively accessible measurements (completed in less than 30 seconds), addressing recent calls to move beyond single-time-point measurements towards sensor-based phenotyping that captures the temporal and environmental dependence of carbon metabolism for breeding applications (Scafaro et al., 2026). By linking repeated SPAD and hyperspectral observations with *PhotoGEA* gas-exchange curve fitting (Lochocki et al., 2025), spectral prediction, biochemical conversion, and a protein-turnover model (Eqs. 1–3), the pipeline converts simple measurements into dynamic physiological traits describing synthesis capacity (*S*_max_), the age-related decline in synthesis (*t*_d_), and degradation (*D*_r_) within the light-harvesting (*N*_c_), electron-transport (*N*_j_), and carboxylation (*N*_v_) pools.

### 4.1 Technical performance and physiological information

The pipeline assumes that rapid hyperspectral measurements, although less precise than direct gas exchange, can recover physiological dynamics when repeated observations are integrated within a mechanistic model. The spectral models showed moderate predictive power with little bias (Fig. S5), supporting their use for comparative inference from temporal trajectories rather than for precise single-point estimation (Roth et al., 2025). Importantly, hyperspectral reflectance resolved photosynthetic capacity beyond what pigment content alone can convey. SPAD constrains chlorophyll and therefore *N*_c_, but the spectral models predicted *V*_cmax_ and *J*_high_, with *J*_high_ subsequently converted to *J*_max_—and hence *N*_v_ and *N*_j_—with dynamics that diverged from those of *N*_c_ (Fig. 6). Had these pools simply co-varied with chlorophyll, reflectance would have been redundant with SPAD; instead, their distinct genotype- and treatment-associated trajectories indicate that reflectance captures carboxylation and electron-transport capacity independently of pigment content. The predictive performance of our PLSR models was slightly lower than in some previous studies, which focused on green leaves measured at a single stage of the leaf lifespan (Meacham-Hensold et al., 2019; Silva-Perez et al., 2018; Zhi et al., 2022). Although our calibration dataset included a similar number of plants, observations spanned almost the full leaf lifespan, from emergence to senescence. Consequently, fewer observations were available within each developmental period, making age-dependent changes in the spectral–physiological relationship more difficult to distinguish from variation in *V*_cmax_ and *J*_high_ within that period. This broader sampling range may reduce predictive precision at individual stages, but it is necessary for resolving photosynthetic capacity throughout leaf development (Calzadilla and Johnson, 2025).

The moderate between-cultivar accuracy also reflects limitations in the physiological information contained in leaf reflectance. PLSR models primarily detect total Rubisco content rather than its activation state and may therefore overestimate functional *V*_cmax_ (Meacham-Hensold et al., 2019; Scafaro et al., 2023). Predicting *J*_high_ was more difficult than predicting *V*_cmax_, partly because fewer *J*_high_ measurements were available and because more proteins contribute to electron transport and its changes over time, which may explain its lower accuracy (Fig. S5). Increasing the calibration set from one to two cultivars produced only sma **l** improvements in leave-one-genotype-out prediction (*V*_cmax_, 0.42 to 0.45; *J*_high_, 0.37 to 0.44, Fig. S8). Although the hyperspectral space of the cultivars used in PLSR calibration covers >99% of the prediction space (Fig. S9), independent validation across broader germplasm remains a priority. Residual variance was homogeneous and did not differ among treatments, but weak thermal-time trends and moderate negative lag-1 autocorrelation remained; *J*_high_ residuals also departed from normality (Supplementary Fig. S10). More *J*_high_ measurements across leaf development may improve prediction.

Together, these validation analyses show that the spectral models recover useful temporal signals across leaf development, but support is stronger for comparative trajectories than for individual measurements or fully independent genotypes (Supplementary Figs. S5–S10).

### 4.2 Practical potential for breeding

The practical value of the pipeline lies in the depth of physiological information obtained from rapid measurements. For the present design, collecting equivalent temporal data by gas exchange would require more than 5,400 h, compared with approximately 30 h of hyperspectral measurements once a valid PLSR model is available. This gain in throughput allows larger germplasm panels to be screened, key stages of leaf development to be captured, and the same plants to be measured repeatedly without destructive sampling. In breeding applications, measurements could target peak capacity, mid-senescence, and late senescence across broad genetic diversity, rather than exhaustively sampling a small number of cultivars (Burnett et al., 2021; Fu et al., 2019; Meacham-Hensold et al., 2019). The resulting traits could also complement genomic selection where physiological phenotyping remains a bottleneck (Furbank et al., 2021; Wang et al., 2026). The persistence of parameter variation across release decades (Supplementary Fig. S20) also suggests that potentially useful diversity remains available for selection. Release-year associations provided no significant evidence of directional breeding change in nitrogen-pool turnover under any treatment (Fig. 7). The weak trends of increasing *S*_max_ and decreasing *D*_r_ in *N*_c_ were non-significant and indicate priorities for future testing rather than clear evidence of directional selection, despite mechanistically agreeing with previously reported breeding-related changes in stay-green traits in the original cultivar panel (Voss-Fels et al., 2019; Wang et al., 2026).

The pipeline is modular: SPAD, hyperspectral prediction, gas-exchange calibration, and curve-fitting components can be updated independently as methods improve. Shared resources such as the Wheat Physiology Predictor could support standardisation and reduce the need for study-specific calibration, although recalibration may remain necessary when new genetic variation enters breeding populations (Furbank et al., 2021; Lamour et al., 2026). Broader adoption would also be supported by transferable hyperspectral models, or by efficient recalibration strategies that minimise the periodic calibration data required (Ji et al., 2024; Wan and Ma, 2024; Xu et al., 2026). Combining the approach with proximal or remote sensing could extend dynamic physiological phenotyping to field experiments (Kronenberg et al., 2021; Roth et al., 2025; Wang et al., 2025). The same framework could also incorporate spectral traits for water status, pigments, and photoprotection, providing a broader description of acclimation under combined stresses (Li et al., 2025).

### 4.3 Biological insights as a proof of concept

The inferred nitrogen dynamics indicate a coordinated sequence of resource reallocation during leaf ageing. Light-harvesting capacity (*N*_c_) began to decline after approximately 400 °Cd, whereas electron transport (*N*_j_) and carboxylation (*N*_v_) persisted until around 600 °Cd, consistent with preferential nitrogen retention in carbon metabolism (Amaral et al., 2024). Figure 6 further shows that acclimation was not coordinated across pools: the treatment ordering of *S*_max_, *t*_d_, and *D*_r_ differed among *N*_c_, *N*_j_, and *N*_v_ . This pool-specific behaviour illustrates the physiological information gained by integrating rapid sensing with the turnover model. The developmental pattern is consistent with the hypothesis that chlorophyll could be reduced without compromising yield under non-limiting light and highlights the potential to optimise canopy nitrogen distribution (Cutolo et al., 2023; Salter et al., 2020; Townsend et al., 2018). Lower chlorophyll in light-saturated upper leaves, combined with greater investment in lower, light-limited leaves, could improve canopy light-use efficiency without a yield penalty (Hikosaka, 2014; Niinemets, 2023). These dynamics may also be relevant to grain protein accumulation. Most wheat grain nitrogen is remobilised from pre-anthesis vegetative stores, and remobilisation is closely linked to senescence (Gregersen et al., 2013; Kong et al., 2016; Masclaux-Daubresse et al., 2010). Our experiment followed the third leaf during the first seven weeks and did not measure grain or yield, so it cannot test the relationship between these dynamics and nitrogen remobilisation directly. The pipeline can instead identify candidate genotypes for targeted follow-up.

The weak cross-environment correlations of the turnover parameters (all r ≤ 0.35, explaining at most 12% of the between-cultivar variance; Figs. S17–S19) show that photosynthetic nitrogen dynamics are highly plastic. What consistency there was fell entirely in the light-harvesting and electron-transport pools, whereas no parameter of the carboxylation pool was conserved across environments. Light-harvesting synthesis capacity may therefore provide a relatively stable reference around which the more plastic carboxylation and electron-transport investments are coordinated to avoid co-limitation (Evans and Clarke, 2019; Quebbeman and Ramirez, 2016; Walker et al., 2014). For breeding, this pool-specific plasticity argues against treating acclimation as a single fixed trait. It also reinforces the need for large, multi-environment screens to identify genotypes with stable or useful responses (Wang et al., 2026, 2025).

### 4.4 Technical innovations, methodological limitations and future development

Two features of the experimental design limit causal interpretation. Treatments were imposed as sequential runs in one chamber and were therefore confounded with run (Supplementary Fig. S2); moreover, a 36.1-h light interruption affected the early FT trajectory (Supplementary Fig. S3). The simulation indicated limited influence on the retained turnover trajectory, but comparisons among treatments remain associational.

Ground-truth errors may originate from both SPAD and gas exchange. The wheat-specific SPAD–chlorophyll relationship used here explained 87% of the variation in measured chlorophyll (Parry et al., 2014), leaving a prediction error with a standard deviation equivalent to approximately 36% of the chlorophyll variation in the calibration dataset. Species -specific calibration therefore remains important, and genotype- and age-related effects require further investigation (Poorter et al., 2023). Leaf structure and environment may also affect SPAD independently of chlorophyll. Nevertheless, the SPAD–chlorophyll relationship remained strong in wheat exposed to salinity and nutrient stress (R² = 0.93; Shah et al., 2017), supporting its use for relative comparisons among treatments.

Estimates of *J*_max_ and *V*_cmax_ also depend on the assumptions used to fit gas-exchange curves (Busch et al., 2024; Duursma, 2015; Lochocki et al., 2025). The variation observed among plants of the same cultivar under identical conditions (Fig. 4A, C) illustrates the intrinsic variability of these measurements and the value of rapid sensing for increasing replication (Furbank et al., 2021; Khan et al., 2021; Silva-Perez et al., 2018). Systematic bias may be more important in senescing leaves, whose low assimilation and limited response to high light or CO₂ can violate the saturation behaviour assumed by standard curve-fitting methods. Predictions at late developmental stages should therefore be interpreted carefully.

A further limitation is the use of fixed biochemical conversion coefficients derived from photosynthetic protein stoichiometry (Buckley et al., 2013) and previously applied in cucumber (Pao et al., 2019a). Although these coefficients should be broadly applicable to C3 plants, protein stoichiometry and catalytic capacity may vary among species and genotypes (McAusland et al., 2020; Sargent et al., 2024), and the coefficients have not been validated specifically for winter wheat. The sensitivity analysis indicated that coefficient uncertainty mainly rescaled *S*_max_ rather than *t*_d_ or *D*_r_, while preserving the comparative conclusions (Supplementary Fig. S12). A systematic wheat-specific offset of these parameters could therefore alter pool magnitudes without changing the relative patterns underlying our conclusions. The consistent positive *S*_max_–*D*_r_ and negative *t*_d_–*D*_r_ correlations (Supplementary Fig. S16) may reflect partial parameter compensation in addition to biological coordination. Together with the lower reproducibility of *D*_r_ across the full panel (Supplementary Fig. S14), this reinforces caution when interpreting individual parameter estimates, and applies with most force to conclusions resting on *t*_d_ or *D*_r_ rather than on *S*_max_.

The bootstrap precision estimates remain conditional on the refitting procedure. The refits were allowed a wider parameter range than the reported model fits, so the resulting intervals describe the information the data carry about each parameter rather than the range the constrained estimator could return. Under these wider bounds, 3.7% of *t*_d_ draws exceeded the upper bound used for the reported fits; no draw was negative. The conclusion-stability result applies only to the biochemical coefficients and should not be generalised to SPAD, PLSR, or leaf-age uncertainty.

Accordingly, the pipeline supports comparative analysis of functional pool dynamics more strongly than precise estimation of nitrogen fluxes. Synthetic recovery supported the same precision hierarchy, with *S*_max_ most identifiable and *D*_r_ least identifiable (Supplementary Fig. S15). Because the synthetic refits were warm-started at the true values and did not repeat the differential-evolution stage, this analysis tests local parameter identifiability under observational noise rather than recovery of the complete fitting procedure from arbitrary starting values. We therefore interpret the inferred parameters as relative differences in functional pool dynamics, not as exact nitrogen fluxes, with *D*_r_ requiring the greatest caution.

### 4.5 Conclusion

This study shows that rapid, accessible measurements can provide more than proxy traits: when combined with physiological modelling and explicit uncertainty assessment, they can capture dynamic features of photosynthetic acclimation across large genetic panels. In 60 winter wheat cultivars and three environments, the pipeline identified pool- and environment-specific trajectories and evaluated hypotheses relevant to breeding. Within the 50-cultivar breeding panel, turnover parameters in the inferred nitrogen pools showed no consistent association with cultivar release year. These identified traits should not yet be treated as breeding recommendations, but as candidates for testing in replicated, multi-environment experiments linked to yield and grain protein. Applying the acclimation-phenotyping pipeline in commercial breeding will require spectral models that accurately track photosynthetic capacity throughout the leaf lifespan, particularly during senescence, together with conversion coefficients validated for wheat. A further step is to model *S*_max_ and *D*_r_ as explicit functions of the microenvironment, allowing the pipeline to describe how the leaf microclimate shapes nitrogen turnover. This would extend the framework from treatment-specific comparison towards a transferable tool for phenotyping acclimation under realistic environmental variability.

## ABBREVIATION MEANING

*S*_MAX,X_: Potential synthesis of nitrogen for the functional pool “x”
*T*_D,X_: Aging factor of the synthesis of nitrogen for the functional pool “x”
*D*_R,X_: Degradation rate of nitrogen in the functional pool “x”
*N*_PH_: Total photosynthetic nitrogen
*N*_X_: Nitrogen invested in the pool “x”: light-harvesting (*N*_c_), electron transport (*N*_j_), and carboxylation (*N*_v_)
*S*_X_: Increase of nitrogen in the functional pool “x”
*D*_X_: Degradation of nitrogen in the functional pool “x”

