## Supplemental Data for "Model-assisted high-throughput phenotyping of photosynthetic acclimation"

Address: Albrecht Thaer-Weg 5, 14195, Berlin, Germany

Supplementary data: 2 tables, 20 figures.

Supplementary Table 1. Cultivars included in the experiment. Cultivar name gives the commercial denomination; registration year is the year of official registration in Germany; country indicates the breeder's country of origin; and breeder identifies the corresponding breeding organisation.

| Cultivar name | Registration year | Country | Breeder |
|---|---|---|---|
| Meister | 2010 | DE | RAGT/Mellinger |
| KWS Santiago | 2011 | GB | KWS Lochow |
| Durin | NA | FR | NA |
| Paroli | 2004 | DE | DSV |
| Patras | 2012 | DE | DSV |
| Anapolis | 2013 | DE | NORDSAAT Saatzuchtgesellschaft |
| Biscay | 2000 | DE | Lochow-Petkus |
| Cubus | 2002 | DE | Lochow-Petkus |
| Forum | 2012 | EU | Nordsaat Saatzucht |
| Potenzial | 2006 | DE | DSV |
| Gaucho | 1993 | USA | USDA-ARS, Oklahoma AES |
| Tarso | 1992 | DE | Saatzucht Hadmersleben |
| Hermann | 2004 | DE | Limagrain-Nickerson |
| Tobak | 2011 | DE | W.v.Borries-Eckendorf |

| | | | |
|---|---|---|---|
| Pionier | 2013 | DE | Deutsche Saatveredelung |
| Kalahari | 2010 | EU | LIMAGRAIN GmbH |
| Intro | 2011 | DE,FR A | RAGT 2N |
| Global | 2009 | DE,AT | RAGT/Mellinger |
| Elixer | 2012 | DE | v. Borris-Eckendorf |
| Inspiration | 2007 | DE | Saatzucht Josef Breun GmbH & Co.KG |
| Edgar | 2010 | DE | LIMAGRAIN GmbH |
| Sponsor | 1994 | FR, IE | Unisigma |
| Impression | 2005 | DE | Saatzucht Schweiger GbR |
| Toronto | 1990 | DE | Strengs Erben |
| Contra | 1990 | DE | Saatzucht Breun |
| Saturn | 1973 | DE | MPI |
| JB Asano | 2008 | DE | Saatzucht Josef Breun GmbH & Co.KG |
| Kerubino | 2004 | DE | Saatzucht Schmid Landau |
| Orestis | 1988 | DE | Strube, Dr. H. |
| Anthus | 2005 | DE | KWS Lochow GmbH |
| Herzog | 1986 | DE | Saatzucht Breun |
| Sorbas | 1985 | DE | Strube, Dr. H. |
| Terrier | 2001 | DE | Nickerson |
| Drifter | 1999 | DE | Nickerson |
| Joss | 1972 | DE | Breustedt |
| Disponent | 1975 | DE | Bayrische Saatzuchtgesellschaft |
| Carisuper | 1975 | DE | Heidenreich und Eger |
| Alidos | 1987 | DE | Saatzucht Hadmersleben |
| Akratos | 2004 | DE | Strube Saatzucht |
| Diplomat | 1966 | DE | Firlbeck |
| Benno | 1973 | DE | Bauer, G. |

| | | | |
|---|---|---|---|
| Caribo | 1968 | DE | Heidenreich und Eger |
| NS 22/92 | NA | SRB | NA |
| Hope | NA | USA | S.Dakota Agricultural Experiment Station |
| Mironovska 808 | 1963 | UKR | Mironovskii institut selektsii i semenovodstva pshenitsy |
| Cordiale | 2003 | GB | KWS UK Limited |
| Premio | 2006 | FR | RAGT |
| BCD 1302/83 | NA | MD | Goertzen Seed Research |
| Camp Remy | 1980 | DE | Unisigma |
| Cajeme 71 | 1971 | MEX | CIMMYT |
| Mexico 3 | NA | MEX | BAZ, Braunschweig Genetic Resources Centre |
| Nimbus | 1975 | DE | Firlbeck |
| Florida | 1986 | USA | NA |
| INTRO 615 | NA | USA | NA |
| Labriego-Inia | 1980 | CHL | INIA Chillan |
| Pegassos | 1994 | EU | Strube Saatzucht |
| Julius | 2008 | NA | KWS Lochow |
| RGT Reform | 2014 | NA | RAGT |
| Nordkap | 2016 | NA | Nordsaat |
| Asory | 2018 | DE | SECOBRA |

Supplementary Table 2. Coefficients used in Equations 5–7 to convert photosynthetic capacity proxies ($J_{max}$, $V_{cmax}$, and [Chl]) into nitrogen content of each functional pool.

| Description | Coefficient | Unit | Value | Reference |
|---|---|---|---|---|
| Chlorophyll per light-harvesting nitrogen | $\chi_c$ | mmol Chl mmol$^{-1}$ N | 0.03384 | Buckley et al., 2013 |
| Chlorophyll per electron-transport nitrogen | $\chi_{cj}$ | mmol Chl mmol$^{-1}$ N | $4.64 \times 10^{-4}$ | |
| Electron-transport capacity per electron-transport nitrogen | $\chi_j$ | µmol $e^-$ mmol$^{-1}$ N s$^{-1}$ | 9.48 | |
| Carboxylation capacity per Rubisco nitrogen | $\chi_v$ | µmol $CO_2$ mmol$^{-1}$ N s$^{-1}$ | 4.49 | |

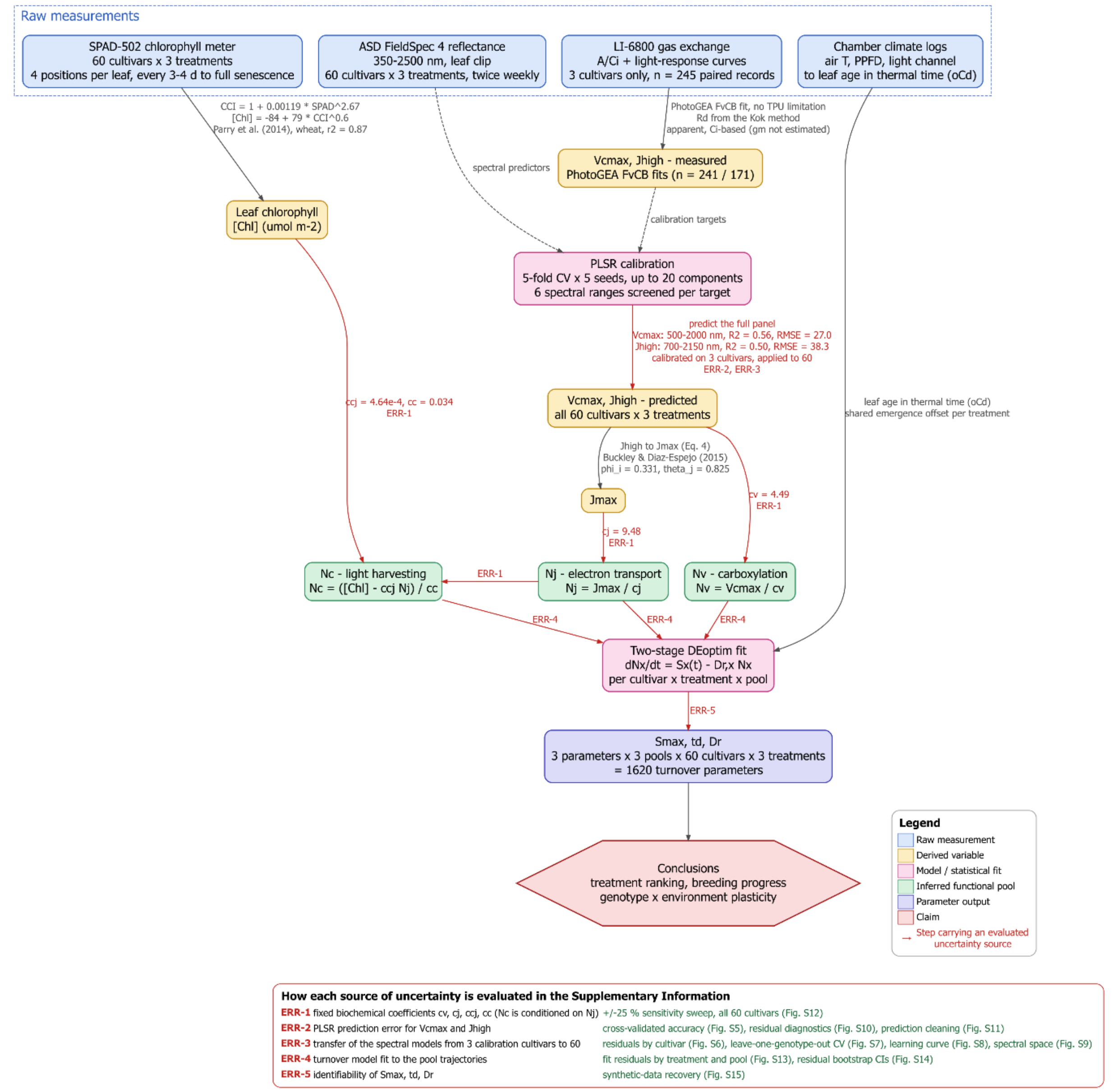


**Supplementary Figure S1.** Computational pipeline and sources of uncertainty. The diagram links four raw measurements, including 1) SPAD-502 chlorophyll, 2) ASD FieldSpec 4 hyperspectral reflectance (350–2500 nm), 3) LI-6800 gas exchange ($A$–$C_i$ curves and light-response curves in three focal cultivars), and 4) chamber climate records, to derived chlorophyll content and estimated or PLSR-predicted maximal carboxylation rate ($V_{cmax}$) and electron transport rate under high light condition ($J_{high}$). $J_{high}$ is converted to $J_{max}$ (Eq. 4 in main text). The derived photosynthetic variables are then converted into photosynthetic nitrogen ($N_v = V_{cmax}/\chi_v$, $N_j = J_{max}/\chi_j$, and $N_c = (\text{Chl} - \chi_{cj} \cdot N_j)/\chi_c$, Eq. 5-7 and Table S2) and fitted with the two-stage *DEoptim* procedure for each genotype × treatment × pool combination. The resulting estimates of parameters $S_{max}$, $t_d$, and $D_r$ (1,620 estimates in total) support analyses of treatment ranking, breeding progress, and genotype × environment plasticity. Node colour indicates raw measurements, derived variables, inferred pools, model fits, parameter outputs, or conclusions. Red edges identify the five sources of uncertainty evaluated in this study: 1) effects from biochemical coefficients, including the dependence of $N_c$ on $N_j$ (ERR-1, Fig. S12); 2) PLSR prediction error (ERR-2, Figs S5, S10, S11); 3) transfer of the spectral models from three to 60 cultivars (ERR-3, Figs S6–S9); 4) fit of the turnover model to the pool trajectories (ERR-4, Figs. S13, S14); and 5) the identifiability and recovery of $S_{max}$, $t_d$ and $D_r$ (*ERR-5*, Fig. S15). The accompanying key links each source to the supplementary figure that evaluates it.

**A**

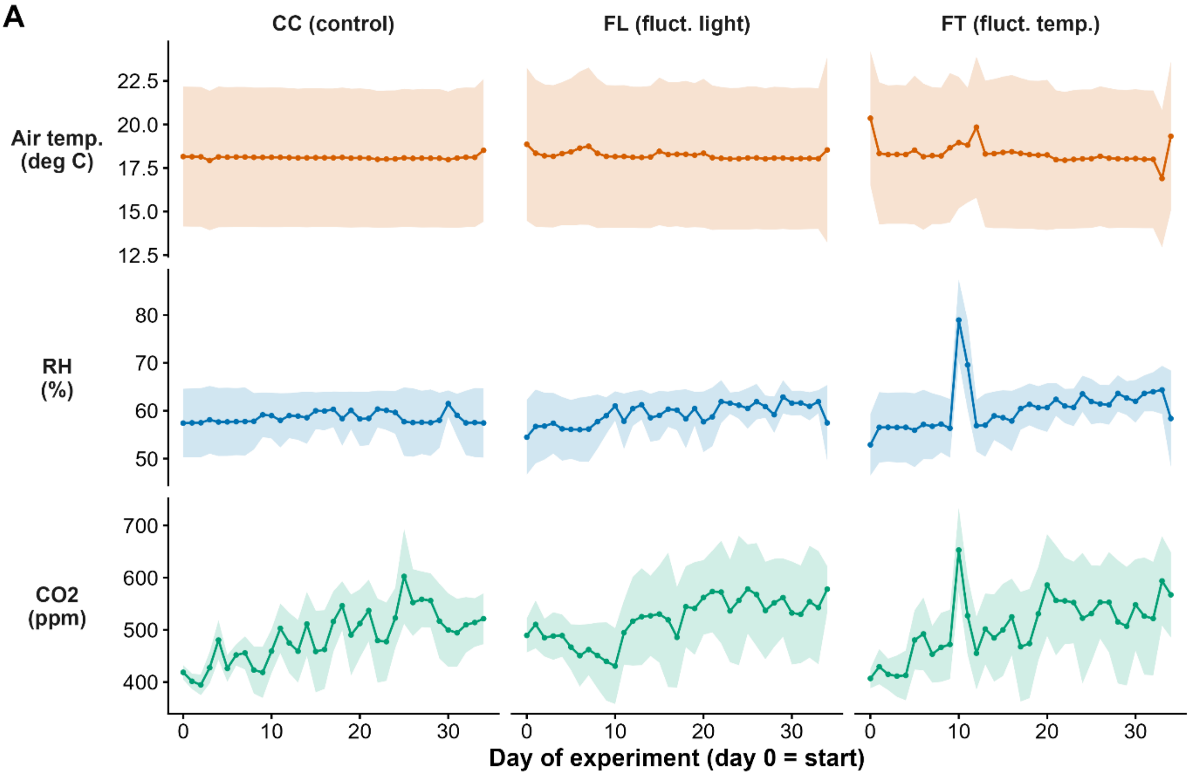


**B**

| Round | Days | Ta (C) | RH (%) | CO2 (ppm) | PPFD (plant) |
|---|---|---|---|---|---|
| CC | 35 | 18.1 +/- 0.1 | 58.5 +/- 1.1 | 487 +/- 49 | 466 [466-466] |
| FL | 35 | 18.2 +/- 0.2 | 59.3 +/- 2.2 | 519 +/- 42 | 442 [366-560] |
| FT | 35 | 18.3 +/- 0.6 | 60.4 +/- 4.6 | 507 +/- 57 | 466 [466-466] |

**Supplementary Figure S2.** Environmental conditions in growth chamber during the experiment. (A) Daily air temperature, relative humidity (RH), and $CO_2$ concentration for CC, FL, and FT. Lines show daily means, and shaded bands show within-day ranges. (B) Mean ± SD conditions over the 35-day monitoring period, including plant-level photosynthetic photon flux density (PPFD). Mean temperature (18.1–18.3 °C), RH (58–60%), and $CO_2$ (487–519 ppm) were similar among runs. The main programmed differences were greater temperature variability under FT (SD = 0.6 versus 0.1 °C under CC) and fluctuating light under FL (mean PPFD = 442 µmol $m^{-2}$ $s^{-1}$; range = 366–560, compared with 466 under CC and FT). Because treatments were imposed sequentially in one chamber, treatment remained confounded with run and contrasts were interpreted as associations. The increasing $CO_2$ baseline reflects sequential operation, while the transient temperature and RH excursion around day 10 of FT coincided with the documented light interruption (Supplementary Fig. S3).

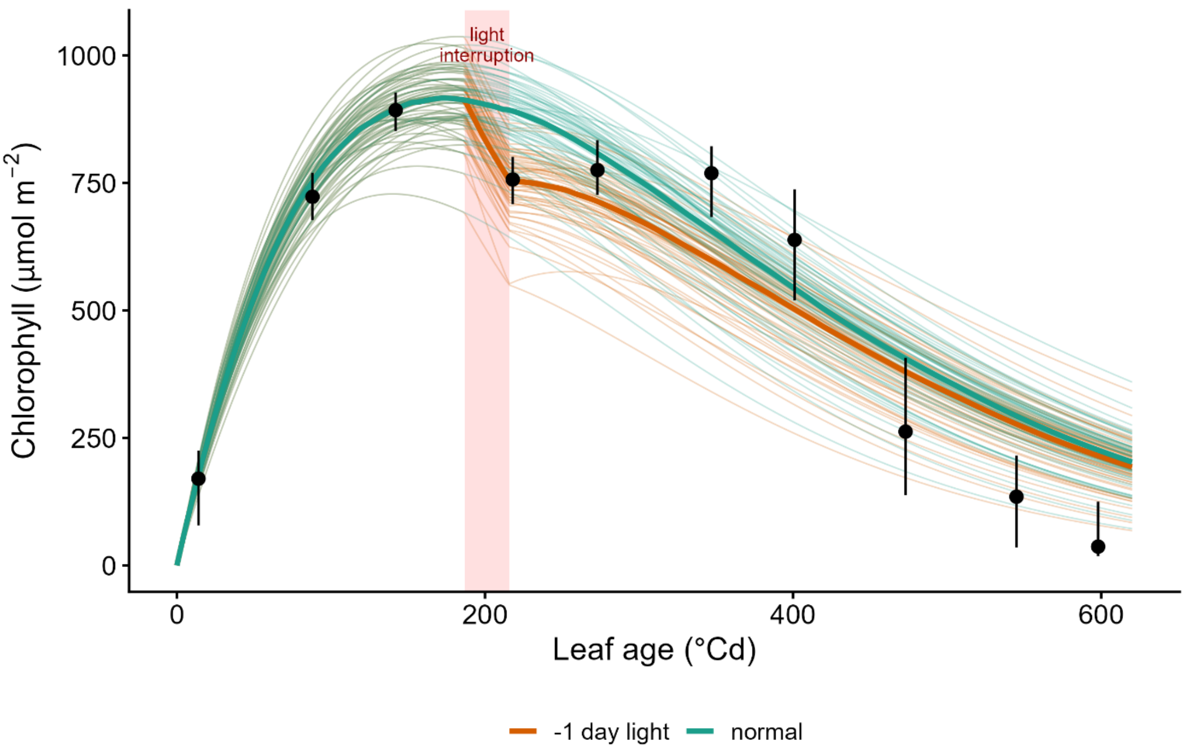


**Supplementary Figure S3.** Observed chlorophyll content under FT (black circles represent median cultivar values and bars the interquartile range across the 60 cultivars) at different leaf ages. Simulated chlorophyll content for each cultivar is shown based on fitted turnover-model with (orange) or without (green lines) one-day light interruption (red interval between 186.9–215.7 °Cd; 28.8 °Cd or 36.1 h). Including the light interruption reduced the model RMSD from 12.9 to 11.3 µmol $m^{-2}$.

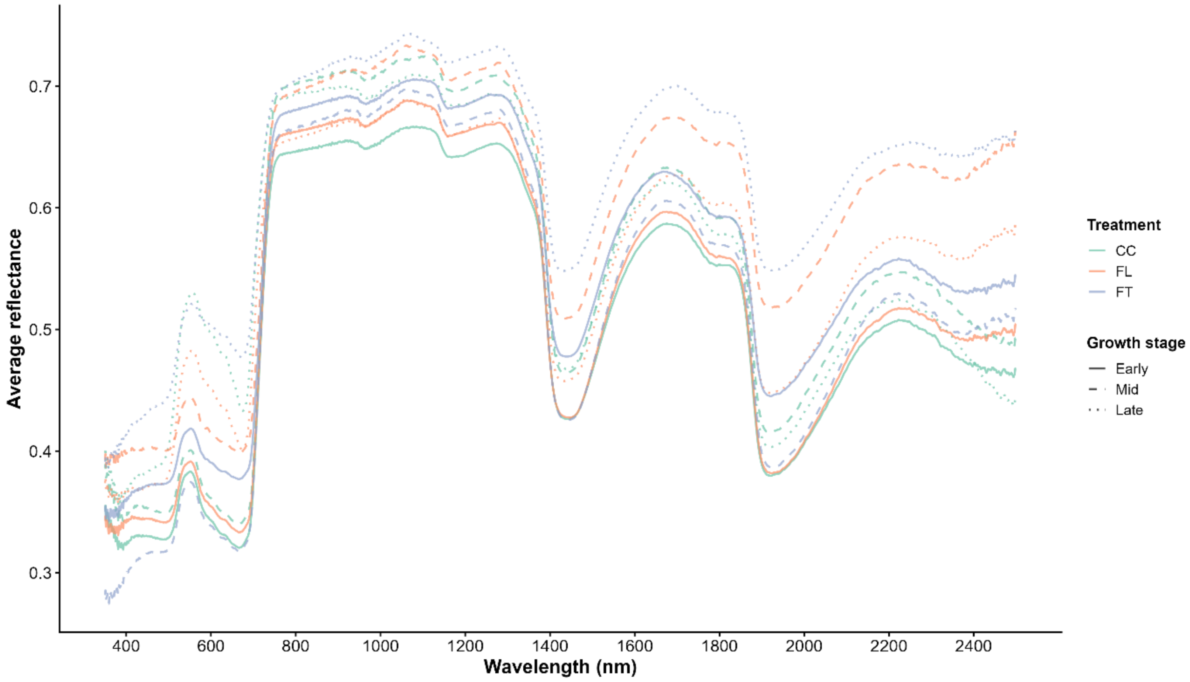


**Supplementary Figure S4.** Mean leaf hyperspectral reflectance (350–2500 nm) across all measurements within each environmental treatment (CC, FL, and FT). The wavelength windows used by the partial least squares regression (PLSR) models were 500–2000 nm for $V_{cmax}$ and 700–2150 nm for $J_{high}$. These windows cover the visible, red-edge, and shortwave-infrared regions associated with pigments, internal leaf structure, water, and protein absorption, while excluding the low-signal extremes of the spectrometer range. The similar mean spectra indicate that the treatments did not produce a large baseline spectral shift; model predictions therefore rely on subtler variation associated with development and genotype.

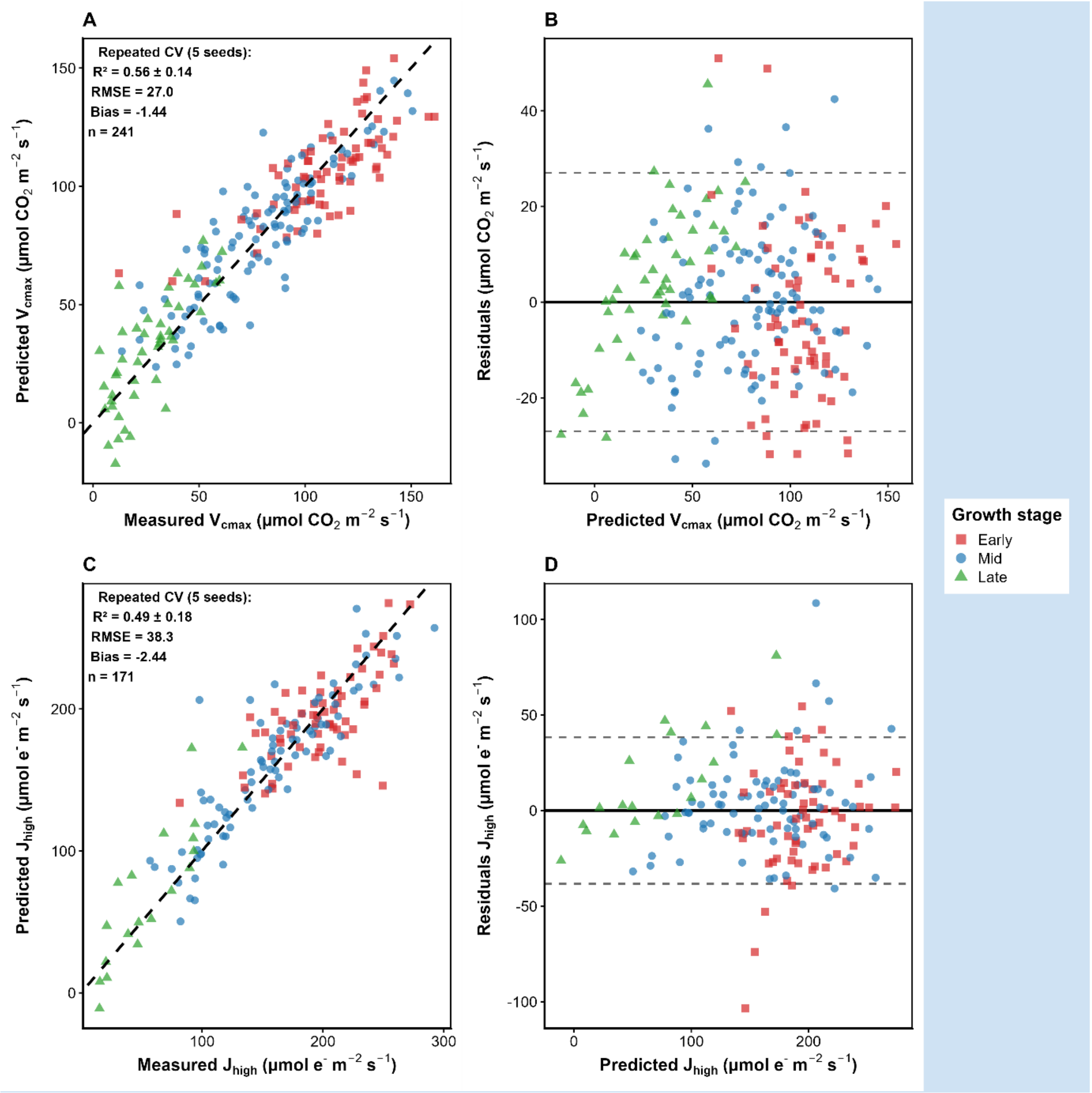


**Supplementary Figure S5.** Validation of the hyperspectral PLSR models for $V_{cmax}$ and $J_{high}$ (ERR-2). (A, C) Comparisons between predictions and gas-exchange measurements; the dashed line shows the 1:1 relationship. (B, D) Comparisons between residuals (predicted minus measured) and predicted values; the solid line marks zero and dashed lines mark ±RMSE. Symbols indicate developmental stage. Validation used five stratified folds with cultivar and 200 °C·d leaf-age classes balanced across folds. Validation repeated five times, and each fold was held out once while the other four were used for fitting PLSR model. $R^2$, RMSE, and bias were calculated from the pooled held-out predictions and reported as mean ± SD across repetitions. The figures show all observations predicted by the final model ($n$ = 241 for $V_{cmax}$ and 171 for $J_{high}$), including observations for model fitting; however, the reported metrics are based only on held-out data. Because plants were measured repeatedly, observations from the same plant on different dates could occur in both calibration and validation sets. Thus, these metrics assess measurement-level predictive performance rather than fully independent plant- or cultivar-level transferability.

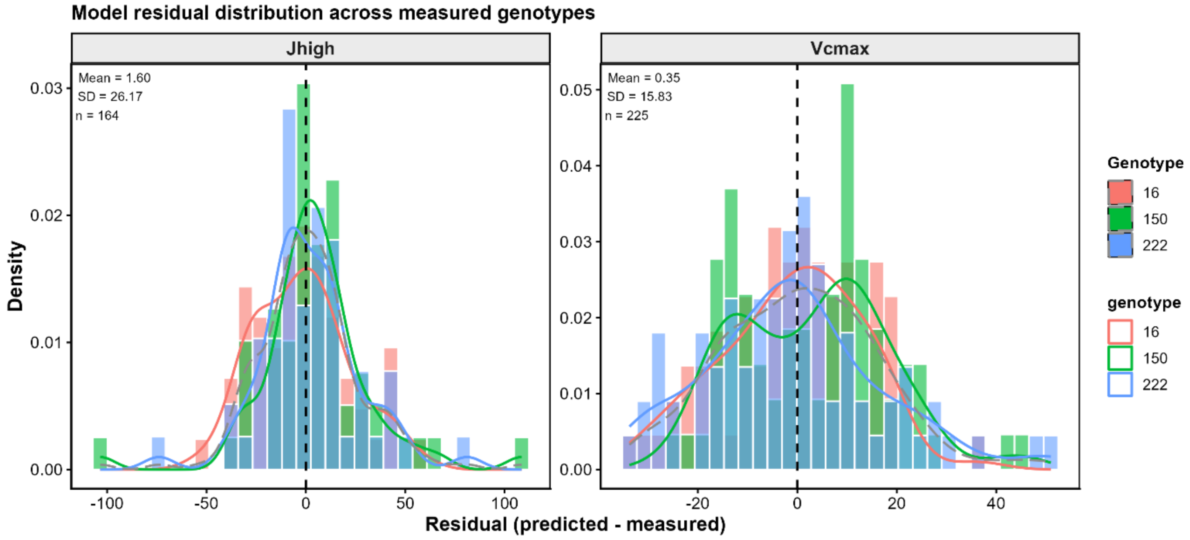


**Supplementary Figure S6.** Distribution of leave-one-genotype-out (LOGO) PLSR residuals (predicted − measured; ERR-3 in Fig. S1), shown separately for each held-out genotype, to assess the transferability of the spectral models across cultivars. In each LOGO iteration, the model was calibrated using two cultivars and evaluated on the third. For $J_{high}$, residuals had a mean of 1.60 and SD of 26.17 (n = 164), whereas for $V_{cmax}$, mean residuals were 0.35 with an SD of 15.83 (n = 225). Mean residuals were close to zero across cultivars, with no pronounced cultivar-specific bias, indicating that the models did not systematically over- or underestimate for individual cultivars. However, the residual spread indicates that substantial prediction uncertainty remains when transferring the models across cultivars. Genotype codes are 222 = RGT-Reform (released in 2014), 150 = Diplomat (released in 1966), and 16 = Durin (exotic; no release year available).

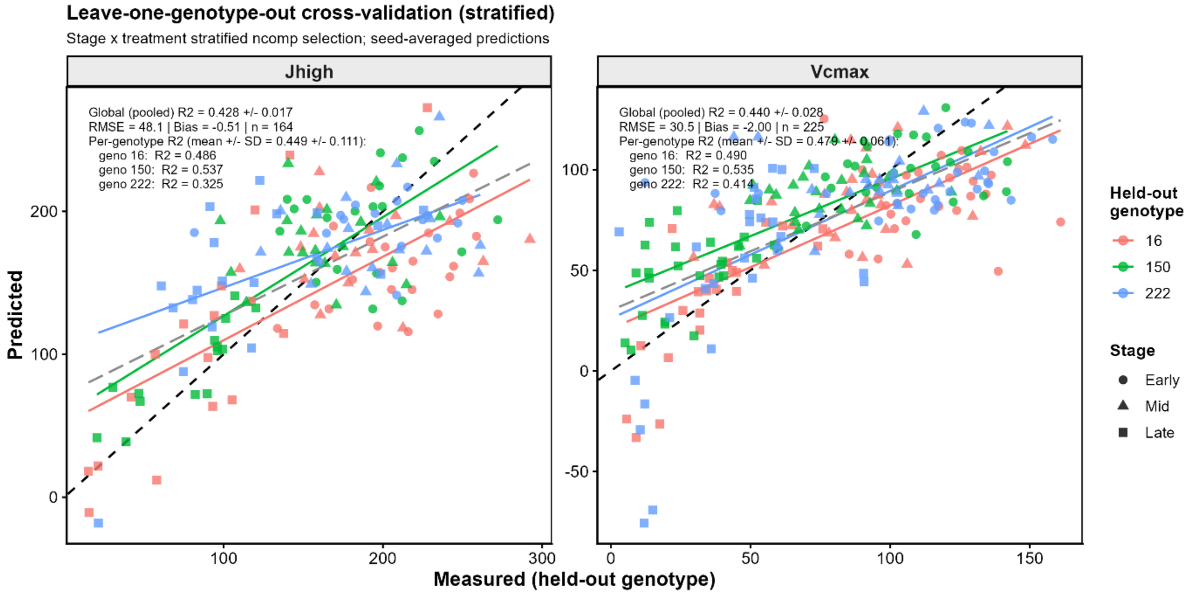


**Supplementary Figure S7.** Evaluation of spectral-model transferability from three to 60 cultivars (ERR-3 in Figs. S1, S6). Leave-one-genotype-out (LOGO) cross-validation was used to assess whether PLSR models calibrated on three cultivars could predict $J_{high}$ and $V_{cmax}$ in previously unseen cultivars. Points represent averaged predictions for each held-out cultivar, coloured by genotype and shaped by developmental stage. The black dashed line denotes the 1:1 relationship, coloured lines show genotype-specific fits, and the grey dashed line shows the pooled fit. For $J_{high}$, the pooled model yielded $R^2$ = 0.428 (RMSE = 48.1, $n$ = 164), and for $V_{cmax}$, $R^2$ = 0.440 (RMSE = 30.5, $n$ = 225). These cross-cultivar predictions provide a conservative assessment of model transferability because each prediction is generated for a cultivar excluded from model calibration.

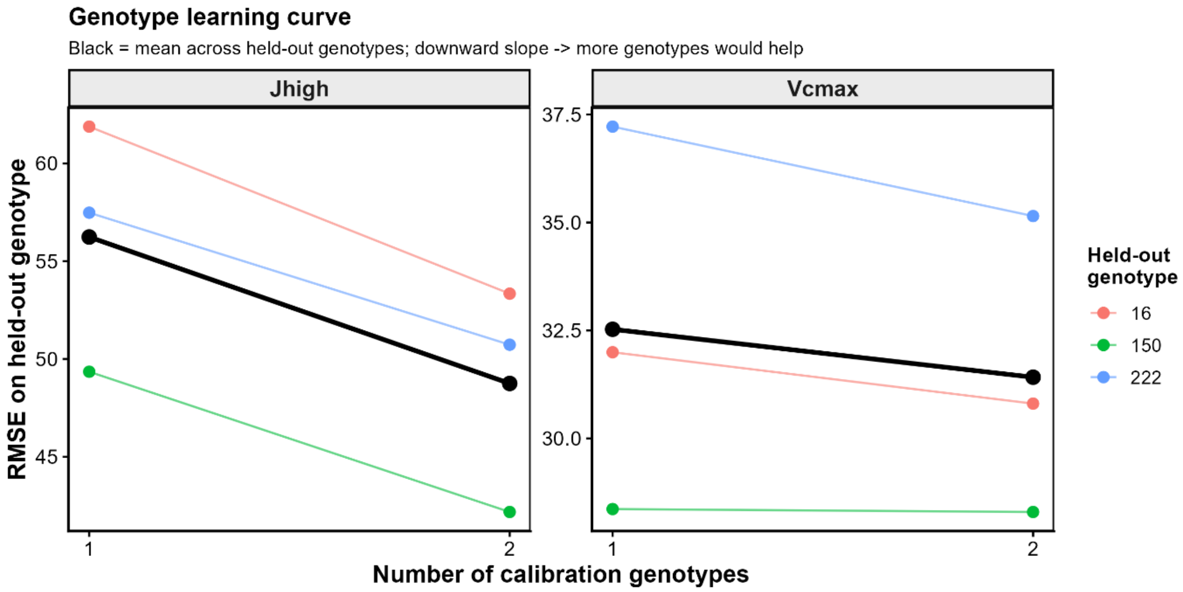


**Supplementary Figure S8.** Effect of calibration-set size on between-cultivar transfer of the spectral models (ERR-3 in Fig. S1). RMSE was compared when predicting a held-out cultivar using either one or two cultivars for calibration. Coloured lines show the mean RMSE for each held-out cultivar across the available calibration combinations (two combinations with one calibration cultivar and one combination with two); the black line shows the mean across all held-out cultivars. Increasing the calibration set from one to two cultivars also doubles the number of calibration observations, so the design cannot distinguish the effect of adding a cultivar from that of adding data. However, RMSE decreased with two-cultivar calibration, particularly for $J_{high}$, indicating improved cross-cultivar transfer with a broader calibration set. The smaller improvement for $V_{cmax}$ suggests that additional calibration cultivars may not improve prediction of this trait.

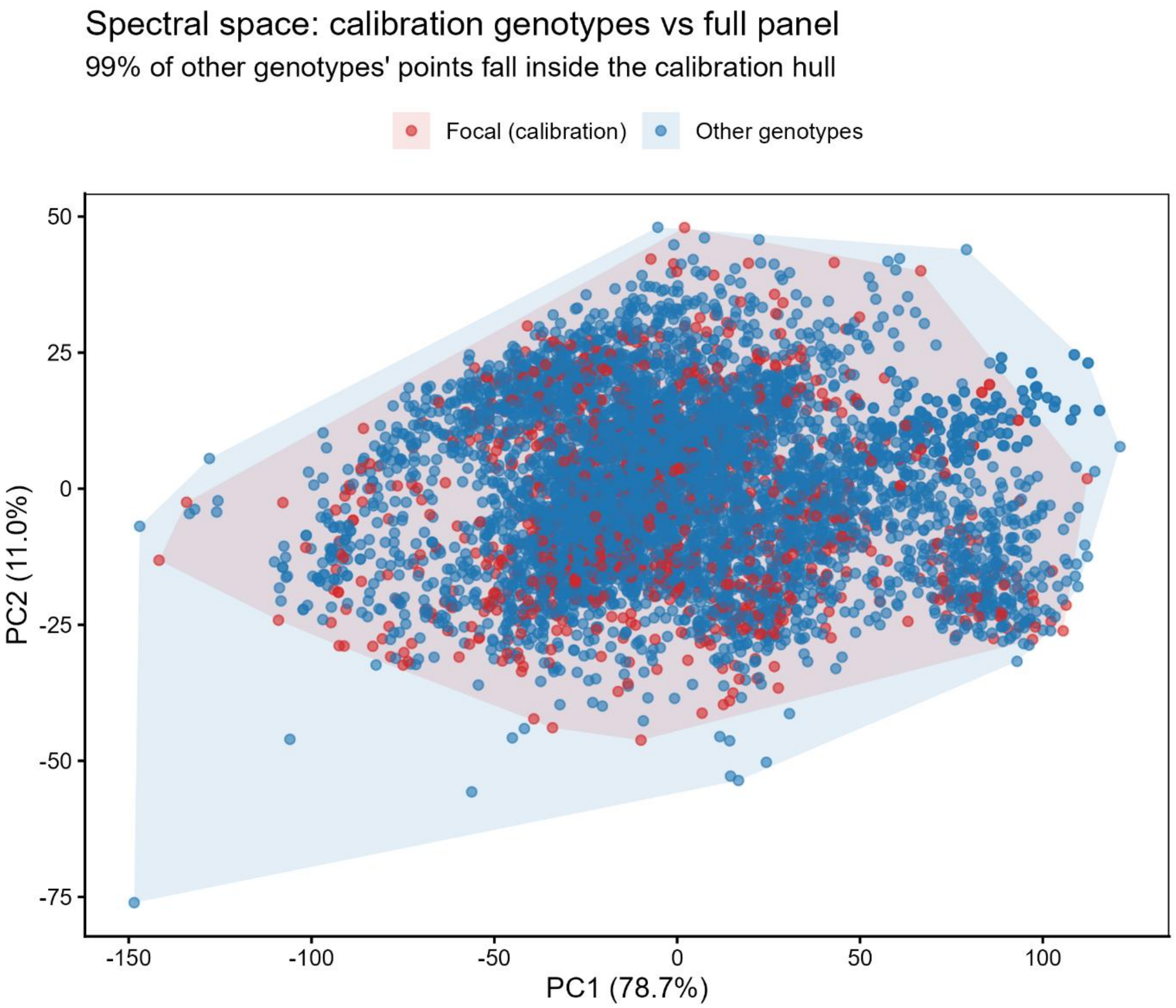


**Supplementary Figure S9.** Evaluation of the spectral domain for model transfer from three calibration to 60 cultivars (ERR-3 in Fig. S1). Principal component analysis (PCA) of leaf reflectance was used to compare the spectral space occupied by the three calibration cultivars (Focal, red points) with that of the remaining 57 cultivars (Other, blue points). Points show individual spectral observations in PC1–PC2 space, with convex hulls indicating the spectral range of each group. Ninety-nine percent of observations from the other cultivars fell within the convex hull defined by the calibration cultivars, demonstrating substantial overlap in the dominant dimensions of spectral variation and providing strong evidence that the calibration set broadly represents the spectral domain of the full 60-cultivar panel. This analysis supports, but does not by itself establish, transferability of the PLSR models to the remaining cultivars.

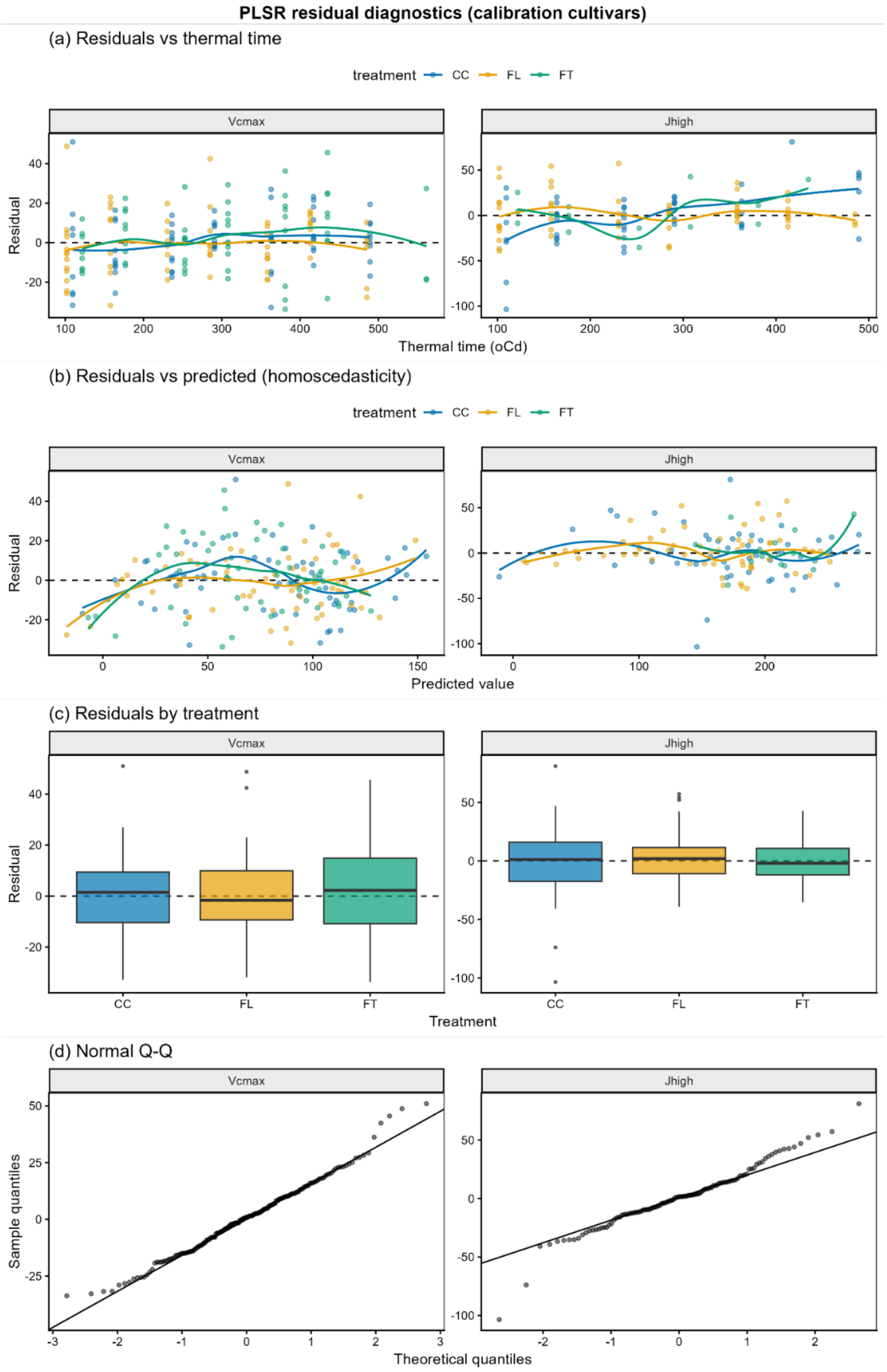


**Supplementary Figure S10.** Residual diagnostics for PLSR models fitted to the calibration cultivars (ERR-2 in Fig. S1). (A) Residuals plotted against leaf age (thermal time); (B) residuals against predicted values; (C) residuals by treatment (CC, FL, and FT); and (D) normal Q–Q plots. Residual variance was homogeneous for both $V_{cmax}$ (Breusch–Pagan $p$ = 0.809) and $J_{high}$ ($p$ = 0.454), with no evidence of treatment-dependent residuals (ANOVA $p$ = 0.523 and 0.864, respectively). Residuals showed weak temporal trends ($p$ = 0.021 for $V_{cmax}$; p = 0.004 for $J_{high}$) and moderate negative lag-1 autocorrelation (−0.41 and −0.40, respectively). $J_{high}$ residuals also deviated from normality (Shapiro–Wilk $p < 0.001$).

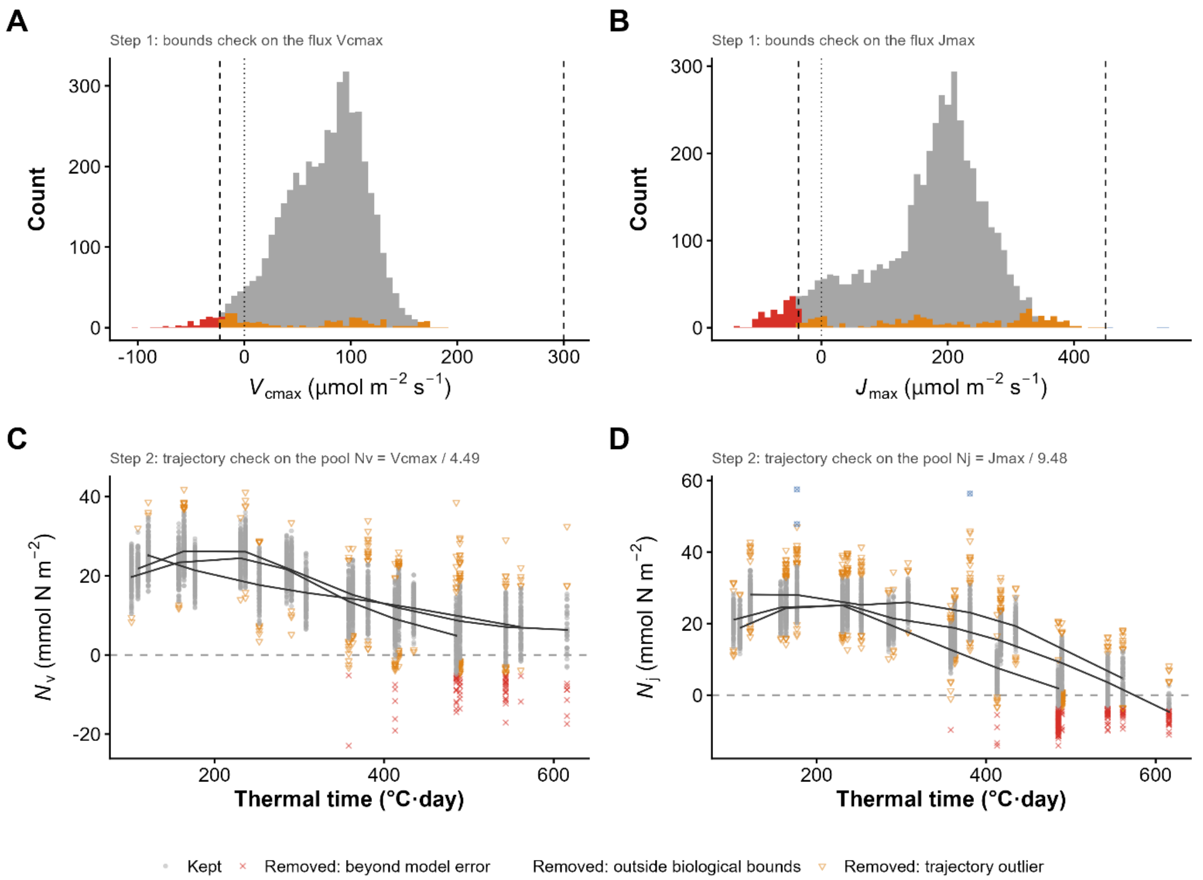


**Supplementary Figure S11.** Outlier detection for PLSR predictions across the 60-cultivar dataset. (A, B) Distributions of predicted $V_{cmax}$ and $J_{high}$, classified as retained (grey), excluded by the tolerance filter (red), or excluded as LOESS trajectory outliers (orange). The tolerance filter excluded predictions below the one-sided negative threshold, defined as 1.5 × half the leave-one-genotype-out (LOGO) RMSE ($V_{cmax}$ < −22.9 µmol $CO_2$ $m^{-2}$ $s^{-1}$; $J_{high}$ < −36.1 µmol $m^{-2}$ $s^{-1}$), or above the realistic physiological maximum; dashed lines indicate these thresholds. (C, D) Corresponding $N_v$ and $N_j$ values across thermal time, with treatment-specific LOESS trajectories shown in grey. The tolerance filter excluded 1.4% of $V_{cmax}$ and 3.8% of $J_{high}$ predictions, primarily during late senescence, while LOESS filtering removed a further 4.5% and 7.0%, respectively, across development. Thus, the filtering procedure was conservative, retaining the large majority of predictions while removing observations inconsistent with physiological limits or the expected temporal trajectories.

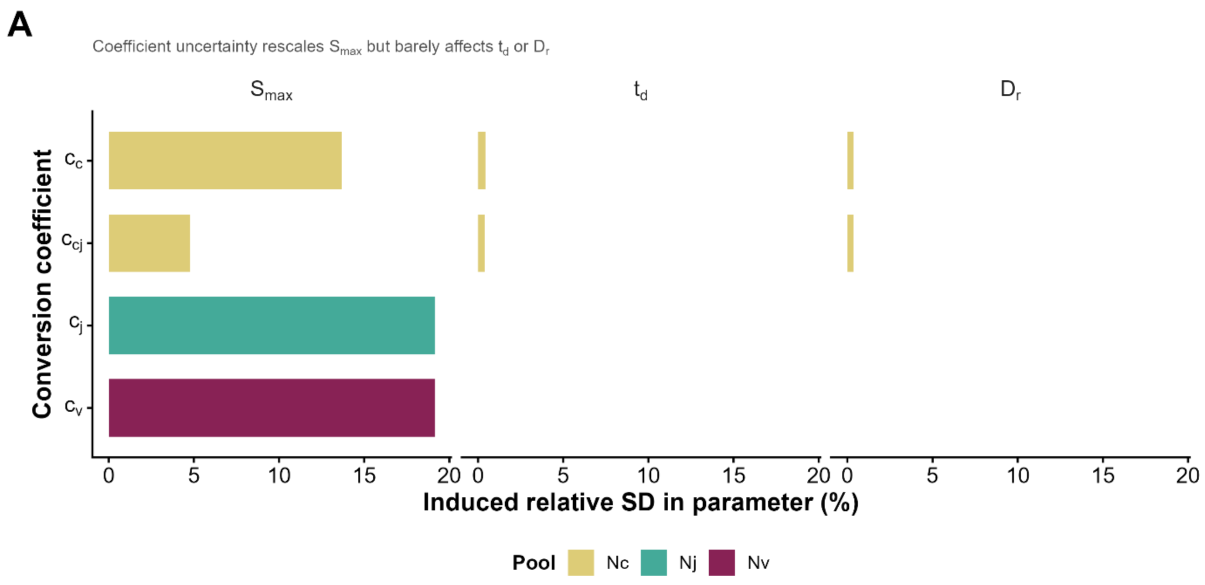


**Supplementary Figure S12.** Robustness of turnover-model parameters to conversion-coefficient uncertainty (ERR-1 in Fig. S1). Sensitivity of $S_{max}$, $t_d$, and $D_r$ to uncertainty in the nitrogen-conversion coefficients ($\chi_c$, $\chi_{cj}$, $\chi_j$, and $\chi_v$). Each of the four coefficients was varied separately at seven levels spanning a conservative ±25% envelope across all 60 cultivars. Bars show the induced relative SD (%) in each parameter, coloured by nitrogen pool ($N_c$, $N_j$, and $N_v$). Coefficient uncertainty primarily rescaled $S_{max}$, which determines absolute pool magnitude, while $t_d$ and $D_r$ changed by <1%. Thus, the shape and timing of the inferred dynamics were more robust than their absolute magnitudes.

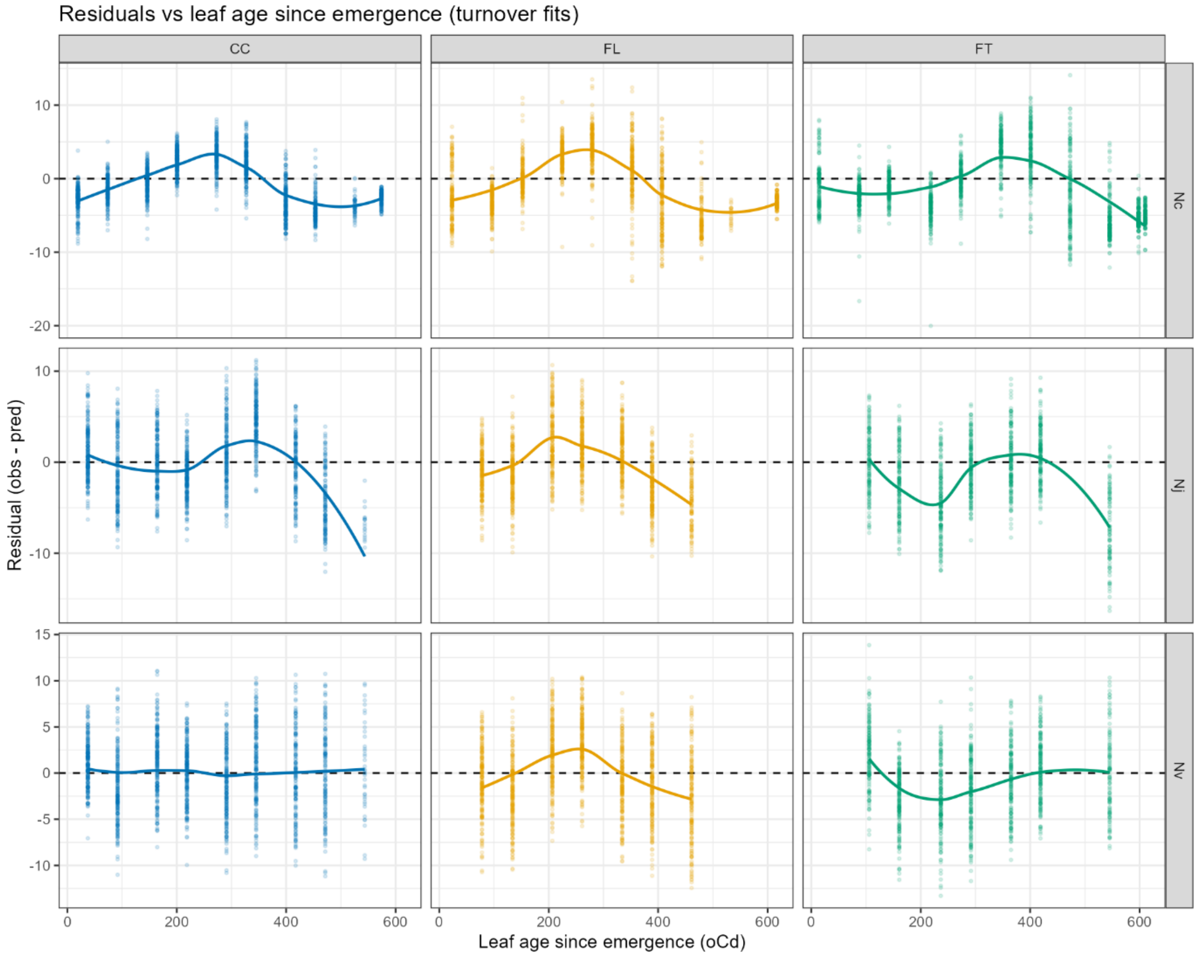


**Supplementary Figure S13.** Turnover-model residuals (observed minus predicted) against leaf age since emergence (ERR-4 in Fig. S1), separated by treatment (columns: CC, FL, and FT) and nitrogen pool (rows: $N_c$, $N_j$, and $N_v$). Points show residuals, coloured lines show LOESS trends, and the dashed line marks zero. Positive residuals indicate model underprediction and negative residuals indicate overprediction. Residuals were broadly centred around zero but showed a general S-shaped temporal pattern: the model tended to overpredict during early development, underpredict around the middle of the trajectory, and overpredict again during senescence. This late-stage overprediction occurred across most treatments and pools, indicating that the model captured the overall developmental trajectories but not all systematic changes in their shape. Under FT, part of the early deviation coincided with the documented light interruption (Supplementary Fig. S3).

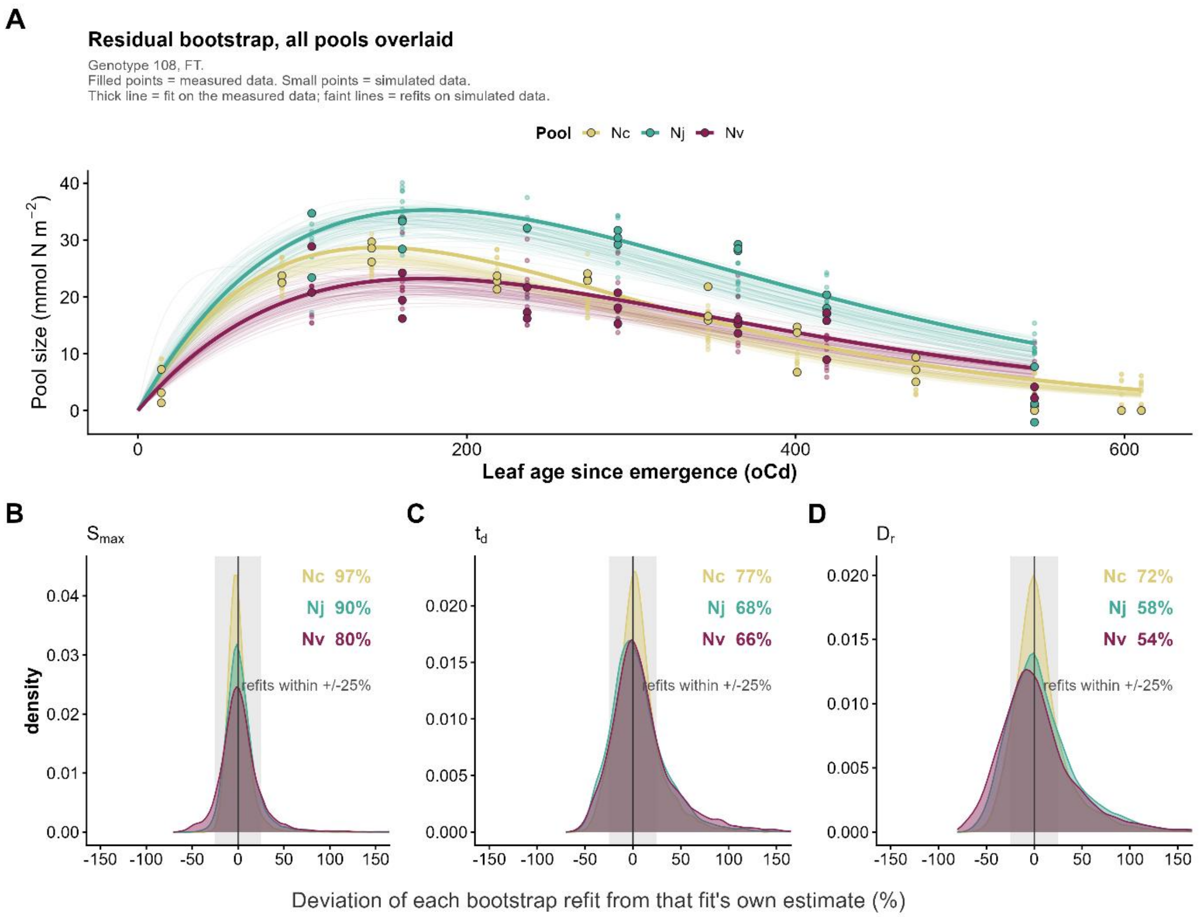


**Supplementary Figure S14.** Residual bootstrap of the three-parameter turnover model (ERR-4 in Fig. S1), shown for a single fit (A) and for every fit in the study (B–D). (A) One example fit: one cultivar × treatment combination (genotype 108, FT), chosen as the condition closest to the panel median in refit reproducibility. Filled points are the measured data; small points are the simulated datasets obtained by adding resampled residuals to the central prediction; the thick line is the fit to the measured data and the faint lines are the 100 refits to the simulated data, for each of the three inferred nitrogen pools. (B–D) All fits: bootstrap distributions of $S_{max}$, $t_d$ and $D_r$ across all 540 cultivar × treatment × pool combinations (60 cultivars × three treatments × three pools, 100 refits each). Each refit is expressed as its percentage deviation from the estimate of the fit it came from, so differences between cultivars are removed by construction and the spread shown is within-fit uncertainty alone; 0% in the x-axis indicates exact recovery of the reported estimate. The shaded band marks ±25%, and the figures give the percentage of refits falling inside it for each pool. Reproducibility was highest for $S_{max}$ (96.6% of refits within ±25% for $N_c$, 90.2% for $N_j$, 80.0% for $N_v$) and lowest for $D_r$ (71.9%, 58.0% and 53.9%), identifying $D_r$—particularly for the carboxylation pool $N_v$—as the least precisely determined parameter. Axes are truncated at ±150%; at most 3.3% of refits for any pool fall outside.

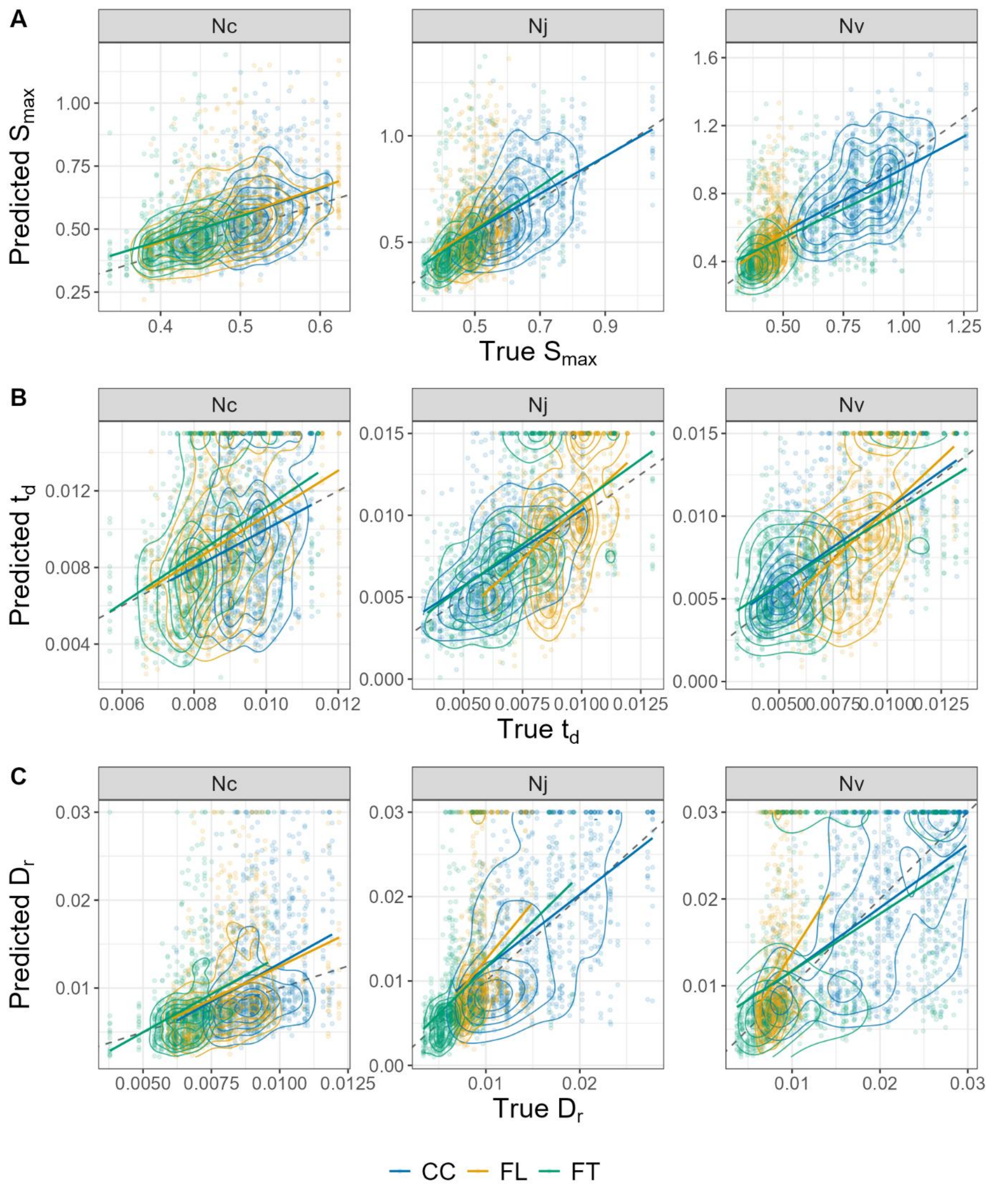


**Supplementary Figure S15.** Parameter identifiability assessed by synthetic-data recovery (ERR-5 in Fig. S1). For every cultivar × treatment × pool combination, the fitted $S_{max}$, $t_d$, and $D_r$ were treated as true values and used to simulate a trajectory at that combination's real measurement times in the emergence frame, with Gaussian noise drawn at the residual standard deviation of the corresponding pool (one noise level per pool, estimated from the residuals of all fits of that pool; the synthetic anchor points used in the original fitting were excluded). Each synthetic trajectory was refitted by bounded Levenberg–Marquardt minimisation warm-started at the true values, using the bounds of the original fits ($S_{max}$: 0.01–2; $t_d$: 0.0001–0.015; $D_r$: 0.001–0.03); the differential-evolution stage of the two-stage procedure was not repeated. Recovered values are plotted against true values for (A) $S_{max}$, (B) $t_d$, and (C) $D_r$, separated by pool and coloured by treatment. Points are individual refits, contours their two-dimensional density per treatment, solid lines the ordinary-least-squares fit of recovered on true values per treatment, and the dashed line the 1:1 relationship. Across 8,100 refits (60 cultivars × 3 treatments × 3 pools × 15 replicates, all converged), recovery was unbiased to slightly positive for all three parameters, but its precision differed markedly between them: $S_{max}$ was recovered most reliably (regression slopes 0.68–1.23; 63% of refits within ±20% of the true value), $t_d$ intermediately (0.81–1.39; 41%), and $D_r$ least reliably (0.66–1.79; 38%), consistent with the low bootstrap precision of $D_r$ (coefficient of variation 26–35%; Supplementary Fig. S14). Fits reaching an optimisation bound—defined as a recovered value equal to the bound to within $10^{-6}$ relative, counted over converged fits—occurred for $t_d$ in 11.4% and $D_r$ in 9.1% of refits, most often for $N_v$ under fluctuating temperature, whereas $S_{max}$ never reached a bound. Bounds were reached because of estimation noise rather than because true values sat at them: for $N_v$ under constant conditions, the median true $D_r$ was 0.016, well below the 0.03 bound.

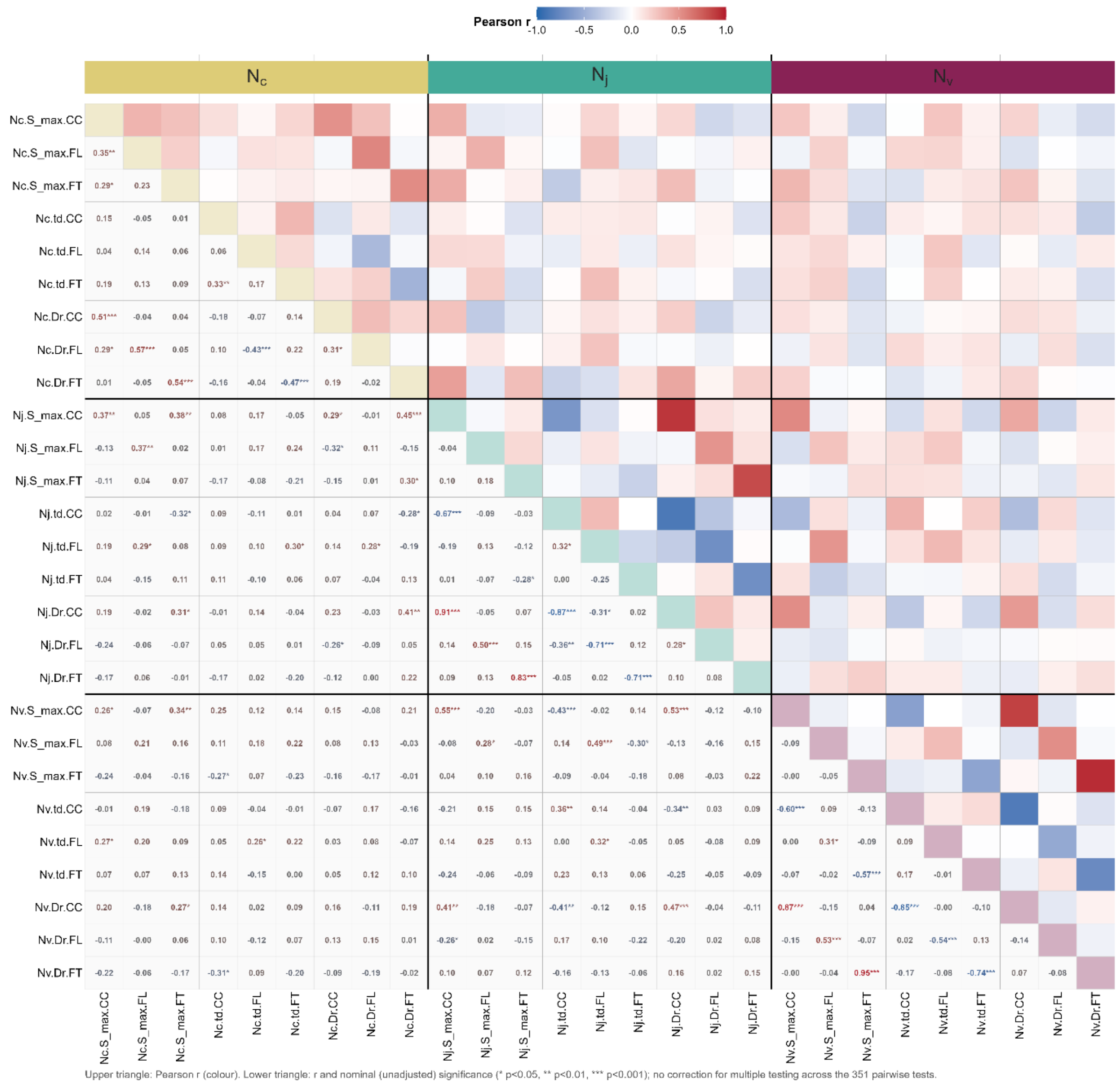


**Supplementary Figure S16.** Correlation matrix of $S_{max}$, $t_d$, and $D_r$ across three nitrogen pools and three treatments (27 variables: pool × parameter × treatment). The upper triangle shows Pearson *r* using the colour scale, and the lower triangle gives the corresponding numerical value. Asterisks indicate raw Pearson significance without multiple-testing correction (*p < 0.05, **p < 0.01, and ***p < 0.001); each correlation used n = 60 cultivars. Thick black lines separate $N_c$, $N_j$, and $N_v$; thin grey lines separate parameters within each pool. Diagonal cells are coloured by pool.

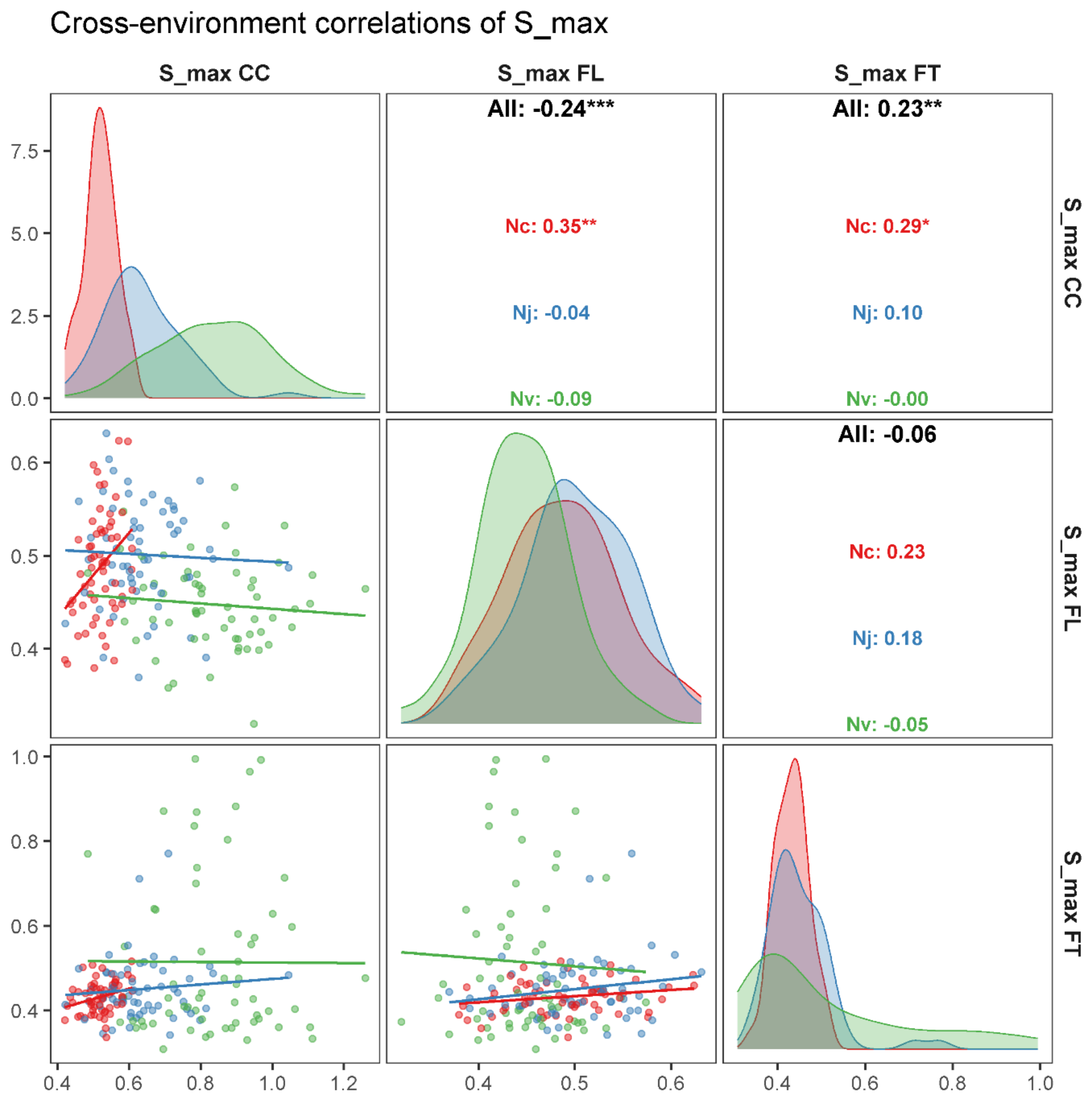


**Supplementary Figure S17.** Cross-environment correlations of $S_{max}$. Panels show pairwise comparisons among CC, FL, and FT. The lower triangle contains scatter plots with linear fits; the upper triangle reports Pearson *r* for all pools combined and for $N_c$, $N_j$, and $N_v$ separately. Diagonal panels show density distributions by pool. Colours indicate $N_c$ (red), $N_j$ (blue), and $N_v$ (green). Asterisks indicate raw Pearson significance without multiple-testing correction (*p < 0.05, **p < 0.01, and ***p < 0.001); each correlation used n = 60 cultivars.

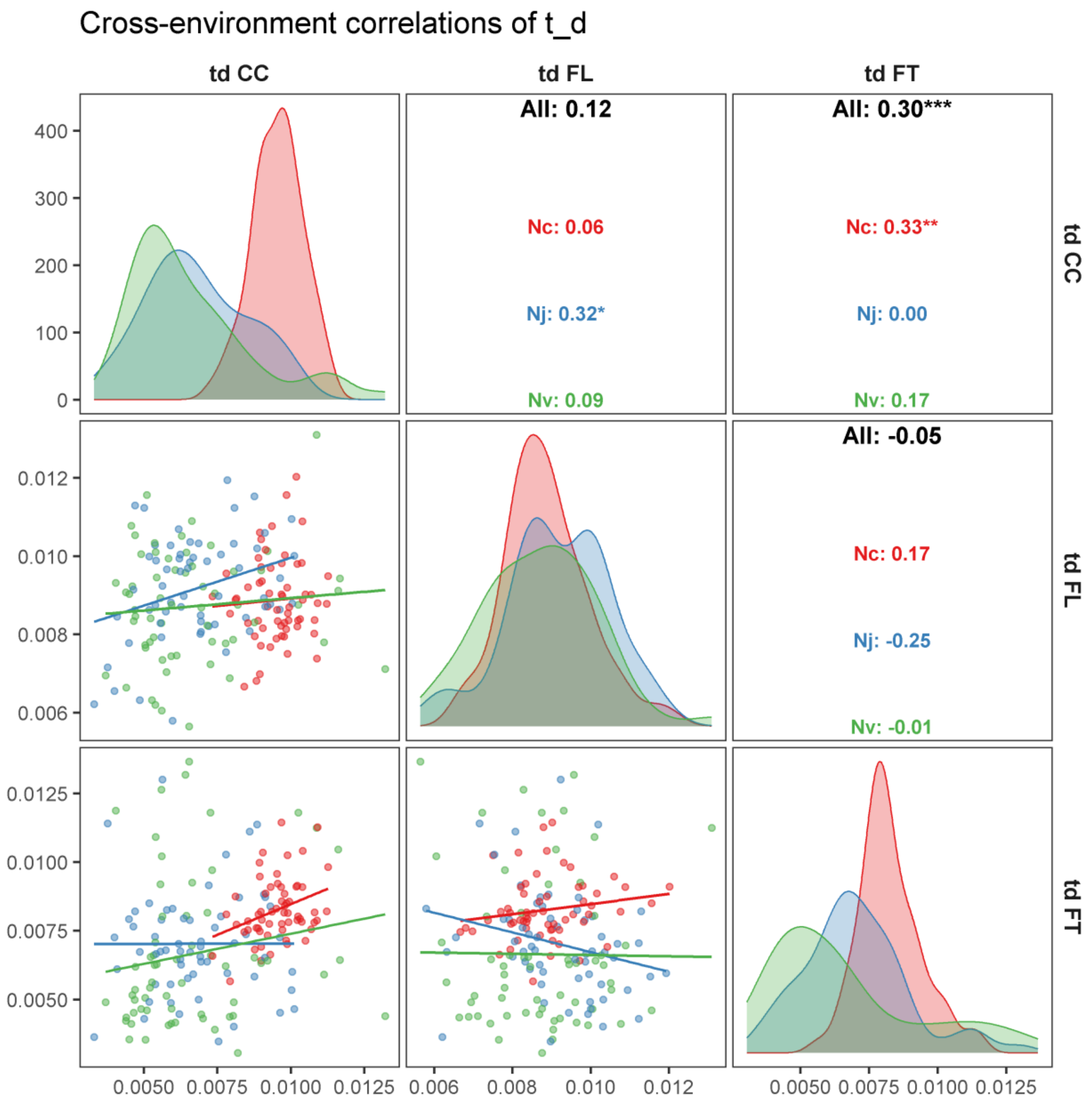


**Supplementary Figure S18.** Cross-environment correlations of $t_d$. Panels show pairwise comparisons among CC, FL, and FT. The lower triangle contains scatter plots with linear fits; the upper triangle reports Pearson *r* for all pools combined and for $N_c$, $N_j$, and $N_v$ separately. Diagonal panels show density distributions by pool. Colours indicate $N_c$ (red), $N_j$ (blue), and $N_v$ (green). Asterisks indicate raw Pearson significance without multiple-testing correction (*p < 0.05, **p < 0.01, and ***p < 0.001); each correlation used n = 60 cultivars.

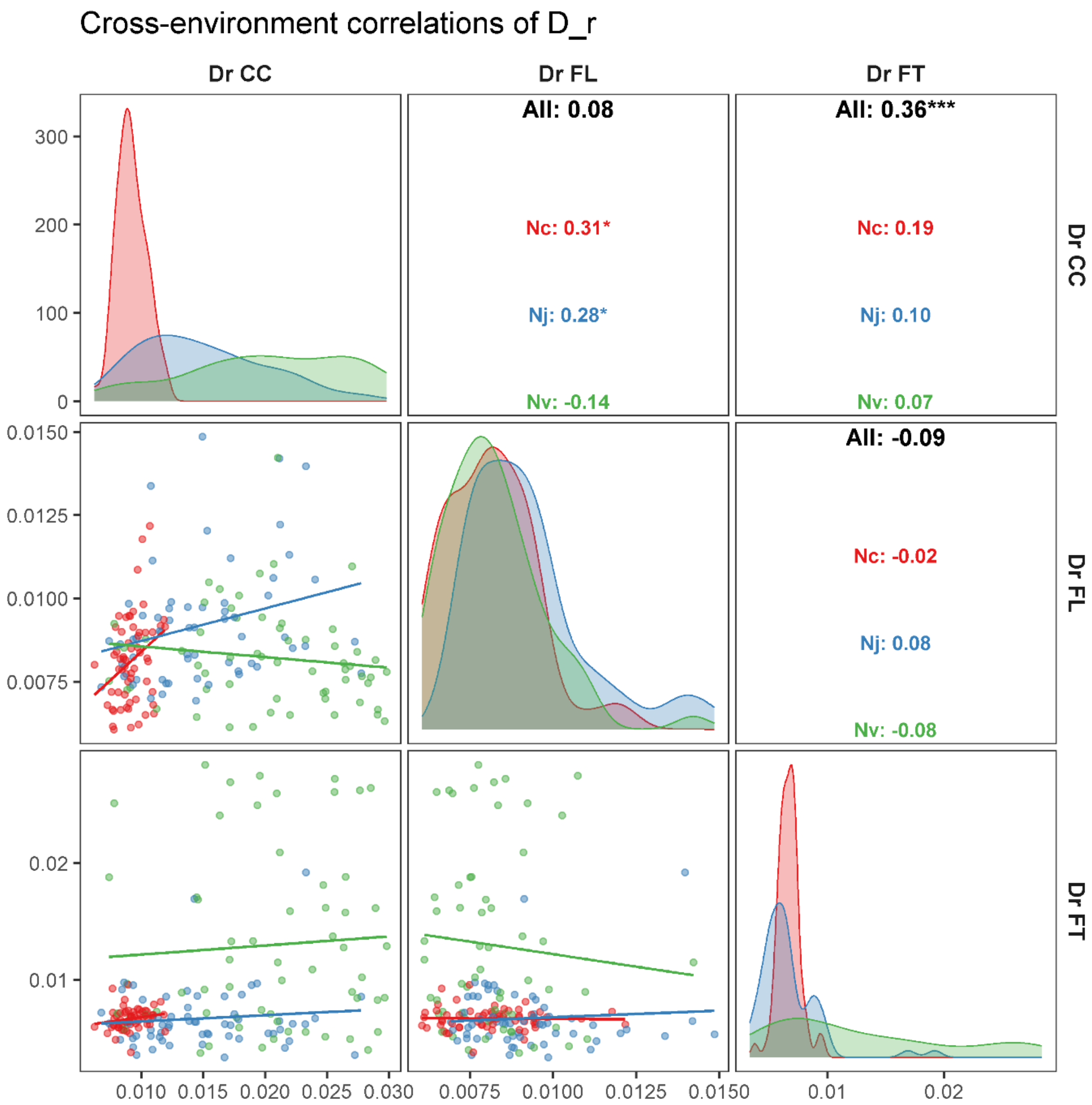


**Supplementary Figure S19.** Cross-environment correlations of $D_r$. Panels show pairwise comparisons among CC, FL, and FT. The lower triangle contains scatter plots with linear fits; the upper triangle reports Pearson *r* for all pools combined and for $N_c$, $N_j$, and $N_v$ separately. Diagonal panels show density distributions by pool. Colours indicate $N_c$ (red), $N_j$ (blue), and $N_v$ (green). Asterisks indicate raw Pearson significance without multiple-testing correction (*p < 0.05, **p < 0.01, and ***p < 0.001); each correlation used n = 60 cultivars.

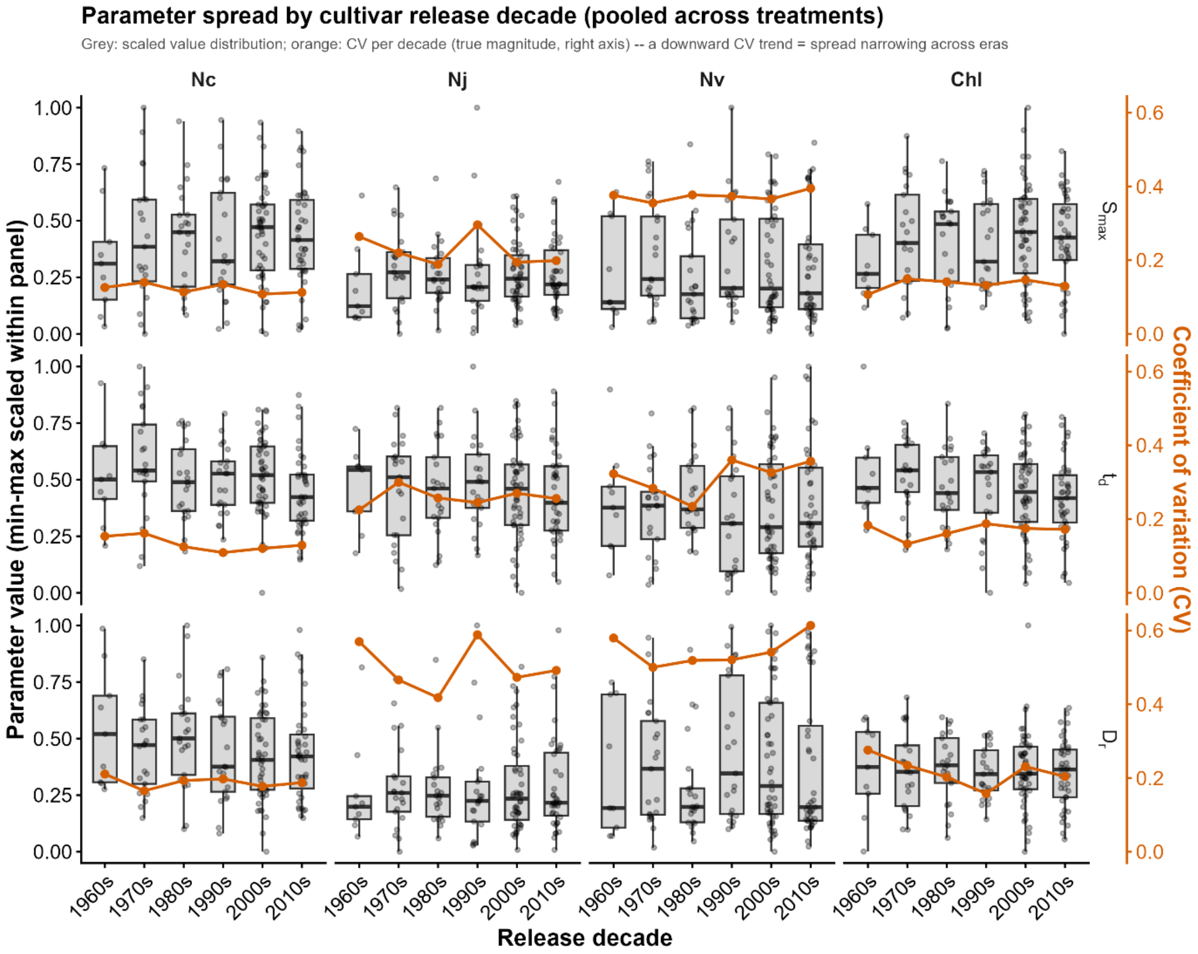


**Supplementary Figure S20.** Turnover-parameter variation across release decades. Cultivars with release-year information (n = 50, excluding exotic accessions) were grouped by decade, with treatment-specific observations pooled within each decade. Columns represent $N_c$, $N_j$, and $N_v$, and rows represent $S_{max}$, $t_d$, and $D_r$. Grey boxplots and points show parameter values min–max scaled within each panel; orange lines show the coefficient of variation for each decade on the right axis. Parameter variation did not narrow systematically in recent decades, indicating no clear convergence in these traits and supporting the absence of strong directional release-year associations (Fig. 7).
